# Breast Cancer Remodels Lymphatics in Sentinel Lymph Nodes

**DOI:** 10.1101/2024.12.30.630756

**Authors:** Dominik Eichin, Akira Takeda, Diana Lehotina, Anni Kauko, Maki Uenaka, Meri Leppänen, Kati Elima, Tapio Lönnberg, Pia Boström, Ilkka Koskivuo, Tero Aittokallio, Maija Hollmén, Sirpa Jalkanen

**Author notes:** **Corresponding author:** Akira Takeda and Sirpa Jalkanen, MediCity University of Turku, Tykistökatu 6, Turku, Finland 20520. Equal contribution (Dominik Eichin, Akira Takeda).

## Abstract

Cancer metastasis to sentinel lymph nodes (LNs) is often the first marker of potential disease progression. Although it is recognized that tumor-induced lymphangiogenesis facilitates metastasis into LNs in murine models, tumor-induced alterations in human lymphatic vessels remain obscure. We used single-cell RNA sequencing to profile lymphatic endothelial cell (LEC) subsets in paired metastatic and non-metastatic LNs obtained from patients with treatment-naïve breast cancer. Tumor metastasis decreases immunoregulatory LEC subsets, such as PD-L1^+^ subcapsular sinus LECs, while inducing an increase in capillary-like CD200^+^ HEY1^+^ LECs. Matrix Gla protein (MGP) was the most upregulated gene in metastatic LN LECs, and its expression on LECs was TGF-β and VEGF dependent. Upregulated MGP promotes cancer cell adhesion to LN lymphatics. Thus, breast cancer cell metastasis to LNs remodels LEC subsets in human LNs and escalates MGP expression, potentially facilitating cancer cell dissemination through the lymphatic system.

## INTRODUCTION

Lymphatics play a critical role in the immune system by efficiently transporting antigens and immune cells into lymph nodes (LNs), which allows timely initiation of immune response or tolerance toward antigens. However, this transportation network can be exploited by cancer cells, contributing to their rapid dissemination. Cancer cells metastasize to draining LNs through lymphatics, colonize LNs and induce immune tolerance against tumor antigens ^1–3^. LN metastasis also protects cancer cells from oxidative stress in subsequent systemic dissemination ^4^. In humans, metastasis of sentinel LNs is a critical parameter for predicting patient mortality ^5–7^. Therefore, understanding the mechanisms by which cancer cells migrate to LNs and promotes metastatic tolerance is of vital importance.

Cancer cells are highly motile, but they also manipulate other cell types to facilitate effective metastasis. One well-known mechanism is tumor-induced lymphangiogenesis. In primary tumors, cancer cells or other cells such as macrophages secrete VEGF-C, which induces the proliferation and sprouting of lymphatic endothelial cells (LECs). This allows cancer cells to metastasize into LNs more effectively ^8–10^. Another lymphangiogenic growth factor, VEGF-D, is secreted from primary tumors and regulates the dilation of collecting lymphatic vessels and subsequent metastasis by regulating prostaglandin generation ^11^. Furthermore, in addition to lymphangiogenesis and the promotion of cancer cell spreading in primary tumors, there is evidence that cancer cells can modulate lymphatic vessels in the draining LNs and thereby facilitate their arrival ^12–14^. Thus, the interaction between LECs and cancer cells is well recognized, but far from deciphered. In addition, many studies have relied on murine models and overlooked LEC heterogeneity, leaving it unclear how cancer cells alter LEC subsets in humans.

Recent studies employing single-cell RNA sequencing (scRNA-seq) have revealed LEC diversity in various organs such as the LN, skin, and intestine ^15–19^. LN LEC subsets are located in distinct areas, such as the subcapsular sinus (SCS) and medullary sinus, and perform subset-specific functions ^20^. LN LECs play crucial roles in regulating immune response through multiple mechanisms, such as controlling immune cell migration, transporting and storing antigens, and presenting antigens to immune cells ^20,21^. Although LEC heterogeneity has been described by us and others in healthy conditions ^15–17^ it has not been studied in detail in diseases such as inflammation and cancer. A recent study using scRNA-seq demonstrated that skin inflammation induces transcriptional changes in SCS floor LECs in mouse LNs ^16^. However, it remains unknown how tumor metastasis to LNs affects LEC subsets in humans and how it impacts cancer metastasis and tumor immunity.

Here we investigated LEC subsets in paired metastatic and non-metastatic LNs from patients with treatment-naïve breast cancer using scRNA-seq. To our knowledge, this is the first report using paired material that allow detection of truly tumor-induced alterations in human LN LECs. We found new LEC subsets accumulating in metastatic LNs and transcriptional changes in established LEC subsets. Matrix Gla protein (MGP) was one of the most upregulated genes in all LN LEC subsets across all the patients. We further confirmed MGP upregulation on LECs in *in vitro* co-cultures with breast cancer cell lines and in the presence of the conditioned medium (CM). We also analyzed the factors behind this upregulation and the consequences of increased MGP on the behavior of LECs.

## RESULTS

### Comparative Single-Cell Analysis of LECs in Human LNs

To understand how cancer cell metastasis impacts LECs in metastatic LNs, we performed scRNA-seq of LECs isolated from metastatic LNs as well as non-metastatic, distant LNs that do not have tumor cells. These samples were obtained from patients with breast cancer undergoing mastectomy with axillary node clearance. LECs were enriched by depleting CD45^+^ cells from single-cell suspensions and subsequently sorting podoplanin (PDPN)^+^ CD31^+^ LECs from distant and metastatic LNs of seven patients with luminal and two with Her2-positive breast cancer (Figure 1a, b, Supplemental Figure 1a). The absence or presence of tumor cells in distant or metastatic LNs was confirmed by staining single-cell suspensions with an anti-pan-cytokeratin antibody (Figure 1a). Pan-cytokeratin positive cancer cells were CD45^-^CD31^-^ PDPN^-^ (Supplemental Figure 1b). PDPN protein expression on LECs, but not on fibroblastic reticular cells (FRCs) or blood endothelial cells (BECs), of metastatic LNs was higher than that in distant LNs, indicating that tumor metastasis alters LECs in metastatic LNs (Figure 1c, Supplemental Figure 1c). The frequency of LECs in LNs varied, but we did not observe proliferation of LECs in the sentinel LNs (Supplemental Figure 1d), as reported in a mouse study ^32^. To integrate scRNA-seq data collected on different days, we employed Seurat version 4, which facilitates the alignment of shared cell populations across diverse datasets and eliminates technical batch effects^22^. Without the alignment and batch correction, cells from different patients were clustered separately (Supplemental Figure 2a). The integrative analysis identified *PROX1*^+^ LEC subsets, *JAM2*^+^ BECs, *COL1A1*^+^ stromal cells, *PTPRC*^+^ leukocytes including *MZB1*^+^ plasmablasts, *KRT19*^+^ cancer cells, and *MKI67*^+^-proliferating cells (Supplemental Figure 2b, c).

**Figure 1.**
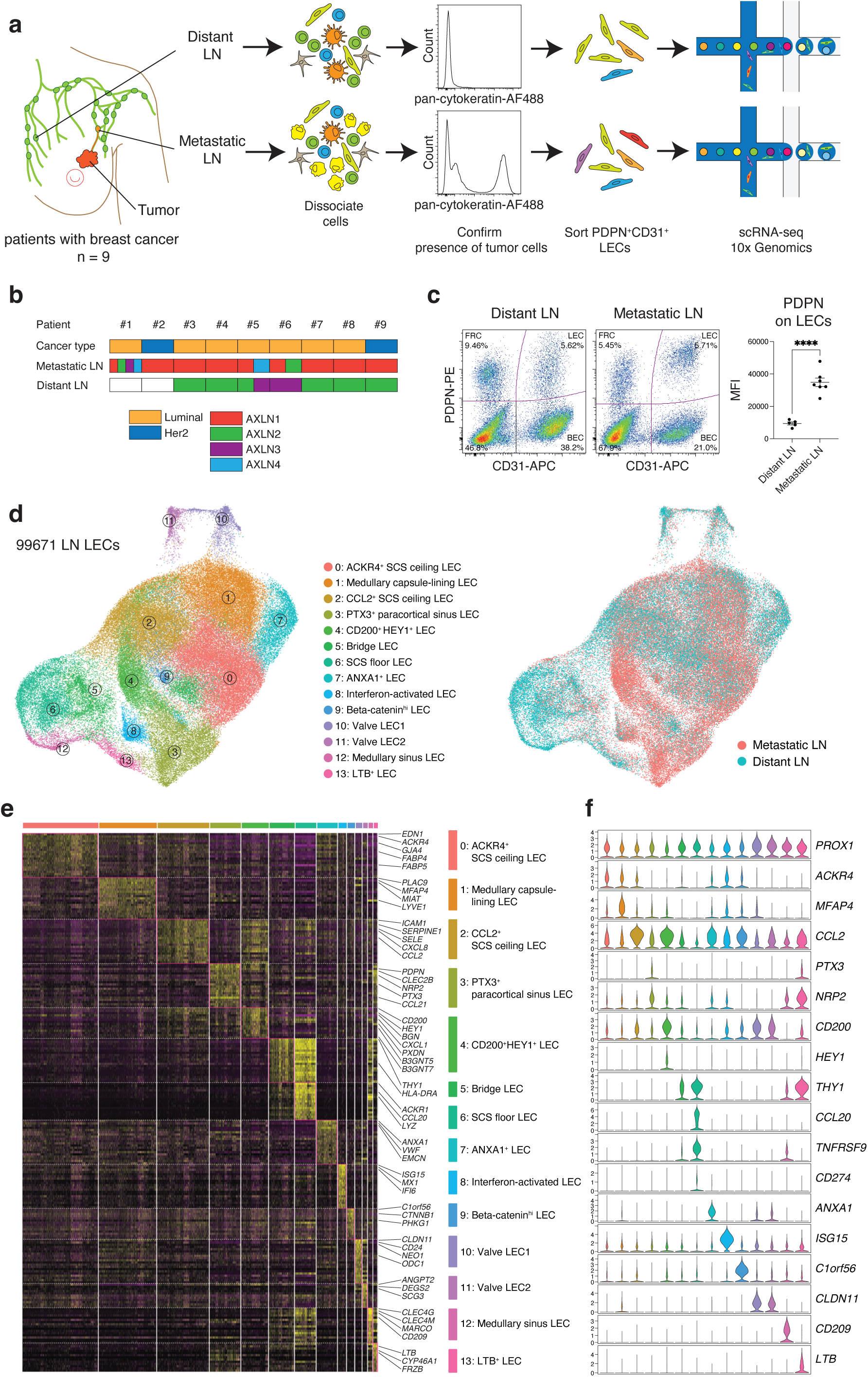
Single-cell analysis of LECs in distant and metastatic LNs from patients with breast cancer. **a,** Workflow. Distant LNs and paired metastatic LNs from nine individuals were used in this study. Cells were dissociated immediately after surgery, and the presence of tumor cells was verified by staining with pan-cytokeratin. CD45^+^ cells were depleted, and PDPN^+^ CD31^+^ LECs were enriched using fluorescence-activated cell sorting. 10x Genomics Chromium scRNA-seq was employed to profile the cells. **b,** Sample information. Patients #2 and #9 had Her2-enriched breast cancer and the others had luminal breast cancer. Paired metastatic and distant axillary LNs were collected from patients #1 to #9. **c,** PDPN and CD31 staining of distant and metastatic LNs. The mean fluorescent intensity (MFI) represents PDPN expression on LECs (mean±SEM, two-tailed, unpaired Student’s t-test), ****p <0.0001. **d,** UMAP plot of 99,671 LECs from distant and metastatic human LNs, color coded by cluster (left) or metastatic state (right). **e,** Heatmap displaying single-cell expression of the top DEGs in LEC subsets, with selected genes labeled. **f,** Gene expression differentiating the 14 LEC clusters, illustrated in violin plots.

To comprehensively characterize LEC subsets, we subclustered *PROX1*^+^ LECs from these enriched populations and analyzed 99,671 LN LECs, including 38,663 LECs from distant LNs and 61,007 LECs from metastatic LNs (Figure 1d). Using unsupervised clustering, we identified 14 clusters within the LEC subsets (Figure 1d). Notably, some LEC subsets, such as cluster 4, were more abundant in metastatic LNs than in distant LNs. To further characterize the LEC subsets in distant and metastatic LNs, we analyzed differentially expressed genes (DEGs) between the LEC subsets (Figure 1e and f). All LEC clusters expressed the typical LEC markers *PROX1* and *FLT4* (also known as VEGFR3) (Figure 1f; Supplemental Figure 2c). Clusters 0–2 highly expressed atypical chemokine receptor 4 (*ACKR4)*, which is selectively expressed by SCS ceiling LECs. Cluster 3 highly expressed pentraxin 3 (*PTX3*), *PDPN*, and neuropilin 2 (*NRP2*), which are markers of paracortical sinuses in human and mouse LNs. Metastasis-induced LEC cluster 4 abundantly expressed biglycan (*BGN*), the transcription factors *HEY1* and *SOX4*, chemokine *CXCL1* and the immunosuppressive molecule *CD200* (Figure 1e and f). Interestingly, this subset selectively expresses acetylglucoaminyltransferase 5 and 7 (B3GNT5 and B3GNT7), indicating a specific glycosylation pattern on this LEC. Clusters 5 and 6 shared specific marker genes, such as *THY1* and *CD74*, but cluster 5 lacked SCS floor LEC markers, including *CCL20*, *TNFRSF9* and *CD274* (also known as PD-L1), which were expressed by cluster 6. Given that bridge LECs (also known as trans-sinusoidal LECs) connect the SCS floor with the ceiling and express both SCS ceiling and floor LEC markers, we assigned clusters 5 and 6 as bridge and SCS floor LECs, respectively. Newly discovered clusters 7, 8, and 9, which were not found in our previous study ^15^, expressed annexin A1 (*ANXA1*), interferon-stimulated gene 15 (*ISG15*), and catenin beta-1 (*CTNNB1*, also known as β-catenin), respectively. Since these clusters also expressed *ACKR4*, they may represent subtypes of SCS ceiling LECs. Annexin A1 is expressed in vascular endothelial cells (ECs) of primary solid tumors, yet *ANXA1*^+^ LECs were found in both distant and metastatic LNs. Clusters 10 and 11 expressed the valve EC marker claudin 11 (*CLDN11)*. Valve LEC1 and LEC2 clusters express neogenin 1 (NEO1) and secretogranin III (SCG3), respectively, and they correspond to LECs on the upstream and downstream sides of valves ^15^. Cluster 12 expressed c-type lectins *CLEC4G*, *CLEC4M*, and *CD209*, which are selectively expressed in LN medullary sinus LECs ^15^. Cluster 13 selectively expressed lymphotoxin-β (*LTB*). Given that this subset expresses significantly both paracortical sinus LEC markers, such as *PTX3* and *NRP2*, and medullary sinus LEC markers including *CLEC4M* and legumain (*LGMN*) (Figure 1e, f), this subset is likely intermediate LECs between medullary and paracortical sinus LECs. This finding aligns with our previous findings, where using LEC trajectory analysis, we found a close relationship between paracortical and medullary sinus LECs ^16^. The newly found LEC subsets in distant LNs were also detected in our published dataset from head and neck LNs by visualizing key marker genes of the subsets (Supplemental Figure 3). Altogether, we demonstrated the highly heterogeneous composition of LECs in human LNs by analyzing a large number of LECs and found significant changes in metastatic LN LECs of patients with breast cancer.

### LN Metastasis Remodels LEC Subsets

Next, we examined changes in LEC subsets within metastatic LNs. The frequency of certain LEC subsets, such as the SCS ceiling and clusters 0, 1, and 2, remained unchanged in metastatic LNs. However, the frequency of other LEC subsets underwent significant changes. Cluster 3 (paracortical sinus LECs) and cluster 4 (CD200^+^ HEY1^+^ LECs) increased in metastatic LNs, whereas cluster 6 (SCS floor LECs), and cluster 12 (medullary sinus LECs) decreased (Figure 2a, b; Supplemental Figure 4a). These alterations in the LEC subsets varied between the patients. However, the trend was consistent across the patients (Supplemental Figure 4b). Changes were observed in patients with both luminal and Her2-enriched breast cancer (Supplemental Figure 4b, c). To analyze differential abundance of cell subsets between distant and metastatic LNs, we used miloR, a scalable statistical framework for differential abundance testing on single-cell datasets^33^ (Figure 2c, d). This analysis revealed a significant enrichment of cluster 4 (CD200^+^HEY1^+^ LECs) and a modest increase of cluster 3 (paracortical sinus LECs) in metastatic LNs. In contrast, there was a significant depletion in cluster 6 (SCS floor LECs) and cluster 12 (medullary sinus LECs) in metastatic LNs. The abundance of other subsets did not change significantly (Figure 2c, d). Moreover, trajectory inference identified six distinct LEC differentiation lineages (Figure 2e). Among these, trajectory 1 (T1) and trajectory (T2) led to cluster 4 (CD200^+^HEY1^+^ LECs) and cluster 6 (SCS floor LECs), respectively, both originating from cluster 2 (CCL2^+^ SCS ceiling LECs). This suggests that LN metastasis preferentially drives LEC differentiation towards T1 lineages rather than T2. SCS floor and medullary sinus LECs highly express inflammatory molecules, such as neutrophil chemo-attractants, compared with other LECs ^15^, and maintain sinus macrophages ^34^, which are important for tumor immunity ^35^. Immunohistochemical analysis of metastatic LNs showed accumulation of cytokeratin^+^ tumor cells within the SCS, cortex near the SCS, and medullary sinuses (Figure 2f). We also observed individual tumor cells traveling through MARCO^+^ medullary sinuses (Figure 2f). Medullary sinuses without tumor cells expressed MARCO, but those containing tumor cells in the same LNs lost MARCO expression, despite maintaining expression of the lymphatic identity marker PROX1 (Figure 2f), indicating that tumor metastasis through the sinuses alters LEC phenotypes. This finding is in line with the decrease of cluster 12 (medullary sinus LECs) in metastatic LNs (Figure 2b-d). We next sought to identify the location of robustly enriched CD200^+^ HEY1^+^ LECs in metastatic LNs (Figure 2g). CD200 was highly expressed in BECs within both metastatic and non-metastatic LNs at comparable levels. However, CD200 expression on PROX1^+^ LECs was significantly elevated in metastatic LNs compared to non-metastatic ones (Figure 2g). Interestingly, capillary-like PROX1^+^ lymphatics expressing CD200 were found near metastatic cancer cells in metastatic LNs, whereas these lymphatics were undetectable in non-metastatic LNs. Moreover, we observed individual cancer cells within capillary-like CD200^+^ lymphatics, suggesting that cancer cells may disseminate through these lymphatics for systemic spread (Figure 2g). Overall, these data show that tumor metastasis through lymphatic sinuses remodels existing LN sinus LECs and generates abnormal LECs, such as capillary-like CD200^+^ HEY1^+^ LECs, and reduces LECs that are important for an immunological response.

**Figure 2.**
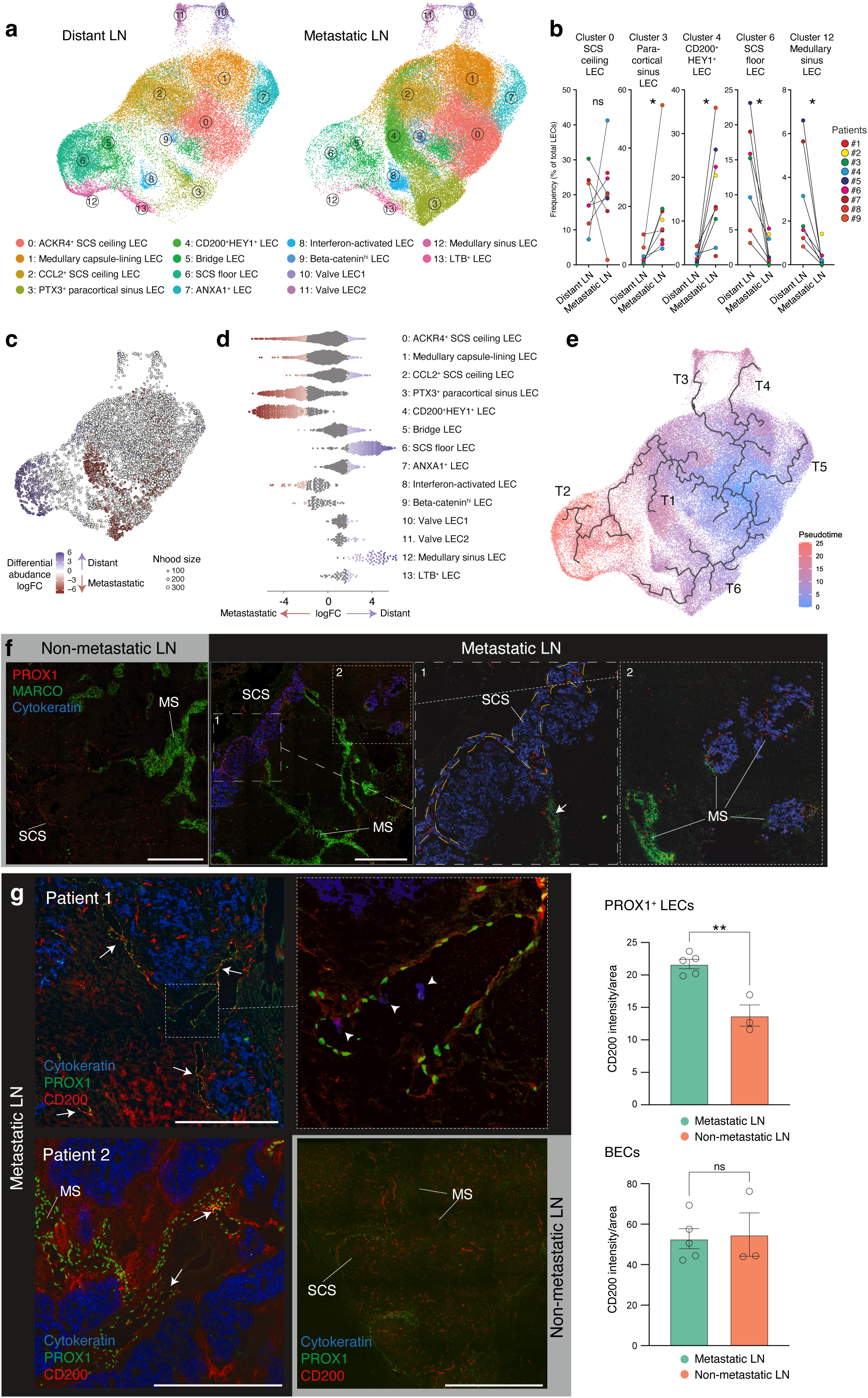
Metastasis through lymphatic sinuses alters LEC subsets in human LNs. **a,** UMAP plot of LECs from distant LNs (left) or metastatic LNs (right), color coded by cluster. **b,** Proportion of each LEC subset in distant and metastatic LNs, calculated from nine patients LECs (two-tailed, paired Student’s t-test), *p <0.05. **c,** Differential abundance testing using miloR. Neighborhoods are colored by their log fold abundance change between distant (blue) and metastatic (red) LNs. Non-differential abundance neighborhoods (FDR > 10%) are colored white. **d,** Beeswarm plot of the cell subset distribution of log fold change between normal and distant LNs. **e,** The trajectories of LN LEC differentiation shown in a UMAP plot. Six distinct LEC trajectories (T1-T6) were identified using Monocle single-cell trajectory analysis. **f,** Immunostaining of cancer cells and LEC markers PROX1 and MARCO in non-metastatic (left) and metastatic LNs (right). Zoomed-in images displaying SCS containing cytokeratin^+^ cancer cells (left) and medullary sinuses (MS) both with and without cancer cells (right). A cancer cell in the MARCO^+^ sinus is indicated by an arrow. Blue, cytokeratin; red, PROX1; green, MARCO. Scale bars, 500 μm. Images are representatives of two individuals with similar results. **g,** Immunostaining of CD200^+^ lymphatics and its quantification in metastatic (upper, black background) and non-metastatic (lower, grey background) LNs. CD200^+^ lymphatics and individual cancer cells within lymphatics are indicated by arrows and arrowheads, respectively. Blue, cytokeratin; green, PROX1; red, CD200. Scale bars, 500 μm. Images are representatives of seven individuals with simar results. SCS: subcapsular sinus; MS: medullary sinus; B: B cell zone. Circles in the bar plots represent biological replicates (mean±SEM, two-tailed, unpaired Student’s t-test). **p <0.005.

### Metastasis-Induced Transcriptional Changes in LN LECs

To investigate transcriptional changes upon tumor metastasis, we identified marker gene expression in neighborhoods corresponding to enriched or depleted LECs in metastatic LNs (Figure 3a, left). This analysis revealed that enriched cells in metastatic LNs express *MGP*, growth arrest specific 6 (*GAS6*), *BGN*, *PDPN*, transcription factor 4 (*TCF4*), and *CD200*. In contrast, depleted cells express inflammatory genes such as *NFKB-IA*, *CXCL1-CXCL3*, *CD74*, and *CD44* (Figure 3a, right). Pseudo-bulk analysis of the scRNA-seq data, irrespective of clusters, also showed the upregulation of *MGP*, *CCL21*, *BGN*, *PDPN*, and microfibril associated protein 2 (*MFAP2*) in metastatic LNs (Figure 3b and c). These genes were upregulated in most LEC subsets in metastatic LNs (Figure 3c, Supplementary Table 1). Immunostaining of MGP revealed its expression on the LN capsule and trabecula but not on LECs in distant LNs, and confirmed its upregulation on PROX1^+^ LECs in metastatic LNs (Figure 3d). Gene enrichment analysis of upregulated genes (log2FC > 0.5, p < 0.05) in metastatic LN LECs revealed a notable enrichment of genes associated with Gene Ontology terms ‘collagen-containing extracellular matrix remodeling’ and ‘external encapsulating structure’, including *MGP*, *BGN*, microfibrillar-associated proteins *MFAP2* and *MFAP4*, and fibronectin 1 (*FN1*) (Figure 3e, left). The most downregulated genes in metastatic LNs included superoxide dismutase 2 (*SOD2*) and the inflammatory chemokines and cytokines *CXCL3*, *IL6*, *CXCL2*, and *CXCL1* (Figure 3a and b). *SOD2* and *CXCL3* downregulation was detected in many clusters (Figure 3c). Gene enrichment analysis of downregulated genes (log2FC < −0.5, p < 0.05) revealed significant enrichment in leukocyte trafficking-associated Gene Ontology terms, including ‘leukocyte cell-cell adhesion’ and ‘leukocyte migration’ (Figure 3e, right). This indicates that proper leukocyte trafficking is impaired in metastatic LNs. Notably, *CD274*, which is expressed by SCS floor LECs, was among the genes downregulated in metastatic LNs. *CD274* expression was not detectable in LN cells, including macrophages and BECs, except for SCS floor LECs (cluster 6) (Supplemental Figure 5).

**Figure 3.**
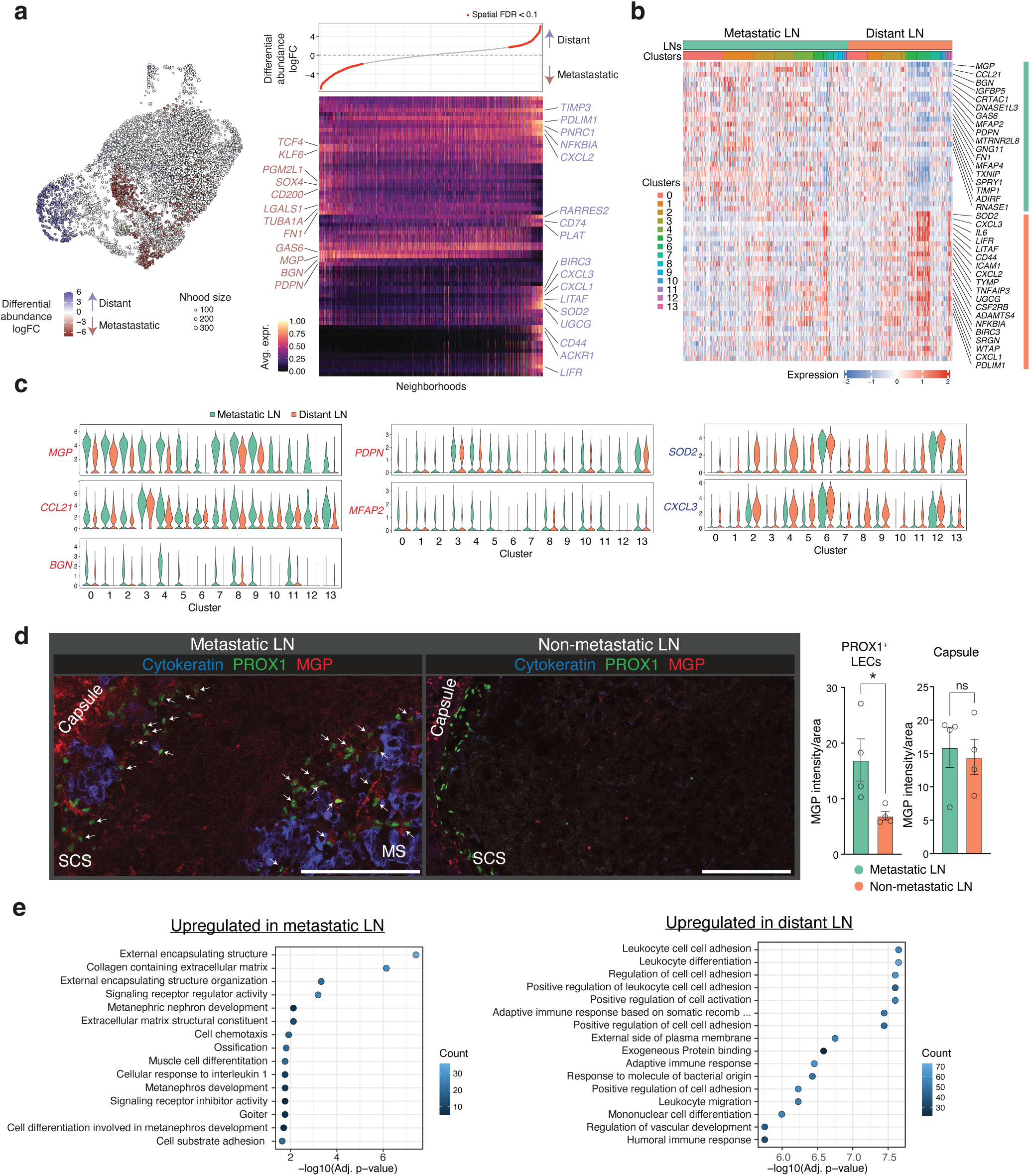
Transcriptional reprograming of metastatic LN LECs. **a,** Differential abundance testing using MiloR and a heatmap of differentially expressed genes between differential abundance neighborhoods in LN LECs. The UMAP plot was shown in Fig. 2c and is inserted here for clarity. In the heatmap, columns and rows represent neighborhoods and differentially expressed genes, respectively. Expression values for each gene are scaled between 0 and 1. The upper panel of the heatmap shows the neighborhood log fold change. FDR, false discovery rate. **b,** Heatmap showing the expression of the top DEGs between metastatic and distant LNs for each cell. Bars above the heatmap indicate the tissue and cluster origin of each cell (LNs, clusters). **c,** Violin plots displaying the top DEG expression between metastatic and distant LNs by cluster, with log-normalized expression value labeled. **d,** Immunostaining of MGP and its quantification in metastatic and distant LNs. Zoomed-in images show SCS and medullary sinuses containing cancer cells and MGP expression on PROX1^+^ LECs (arrows). Blue, cytokeratin; red, MGP; green, PROX1. Scale bars, 125 μm. Images are representative of four individuals with similar results. Circles in the bar plots represent biological replicates (mean±SEM, two-tailed, unpaired Student’s t-test). *p <0.05. **e,** Gene Ontology (GO) enrichment analysis of the top DEGs between metastatic and distant LNs using EnrichR package.

### NicheNet intercellular communication analysis predicts the mechanisms of lymphatic remodeling in metastatic LNs

To understand the mechanisms by which LN lymphatics are remodeled upon LN metastasis, we performed NicheNet analyses. This method predicts the link between ligands from sender cells and changes in gene expression in the receiver cells using prior knowledge on signaling and gene regulator networks^36^. In addition to LECs, metastatic and distant LNs contained lymphocytes, macrophages, plasmablasts, cancer cells, non-endothelial stromal and BECs (Figure 4a). We applied NicheNet to predict which ligand-receptor interactions could drive the DEGs found in CD200^+^ HEY1^+^ LECs and SCS floor LECs, both of which were mostly affected in metastatic LNs. Thus, we designated CD200^+^ HEY1^+^ LECs (cluster 4) or SCS floor LECs (cluster 6) as receivers and the other cell types in LNs as senders that express specific ligands to alter gene expression in the receivers (Figure 4b). *MGP, PDPN, FN1, HEY1 and SOX4*, which are highly detected in enriched LECs in metastatic LNs (Figure 3a), were among the predicted target genes in CD200^+^ HEY1^+^ LECs. The top predicted ligands that induce signatures of CD200^+^ HEY1^+^ LECs included TGF-β, VEGF-A and VEGF-C (Figure 4c, d). TGF-β was commonly expressed by multiple cell types (Figure 4d), including cancer cells, stromal cells and BECs, and its expression was upregulated in metastatic LNs (Figure 4e), potentially driving the expression of *MGP*, *FN1*, *SOX4*, *PDPN* and *HEY1* in LECs (Figure 4c). A transcription factor SOX4 is induced by TGF-β ^37,38^. VEGF-A is highly expressed by macrophages (Figure 4d), and may induce *MGP* and atypical chemokine receptor *ACKR3,* which is a receptor of CXCL12 (Figure 4c). CD200^+^ HEY1^+^ LECs express TGF-β receptors TGFBR2 and TGFBR3, and integrin alpha v ITGAV and integrin beta 1 ITGB1, which can activate a latent form of TGF-β, as well as receptors for VEGF-A and VEGF-C. Notably, TGF-β-dependent LRRC15^+^ cancer-associated fibroblasts ^39^ were only found in metastatic LNs, indicating that TGF-β signaling is upregulated in metastatic, but not in distant LNs (Supplemental Figure 6). In contrast, the top predicted ligands that induce signatures of SCS floor LECs in distant LNs included insulin like growth factor 1 IGF1, Epstein-Barr virus induced 3 EBI3 and LTB, which were mainly expressed by LN stromal cells, macrophages, and lymphocytes, respectively (Supplemental Figure 7).

**Figure 4.**
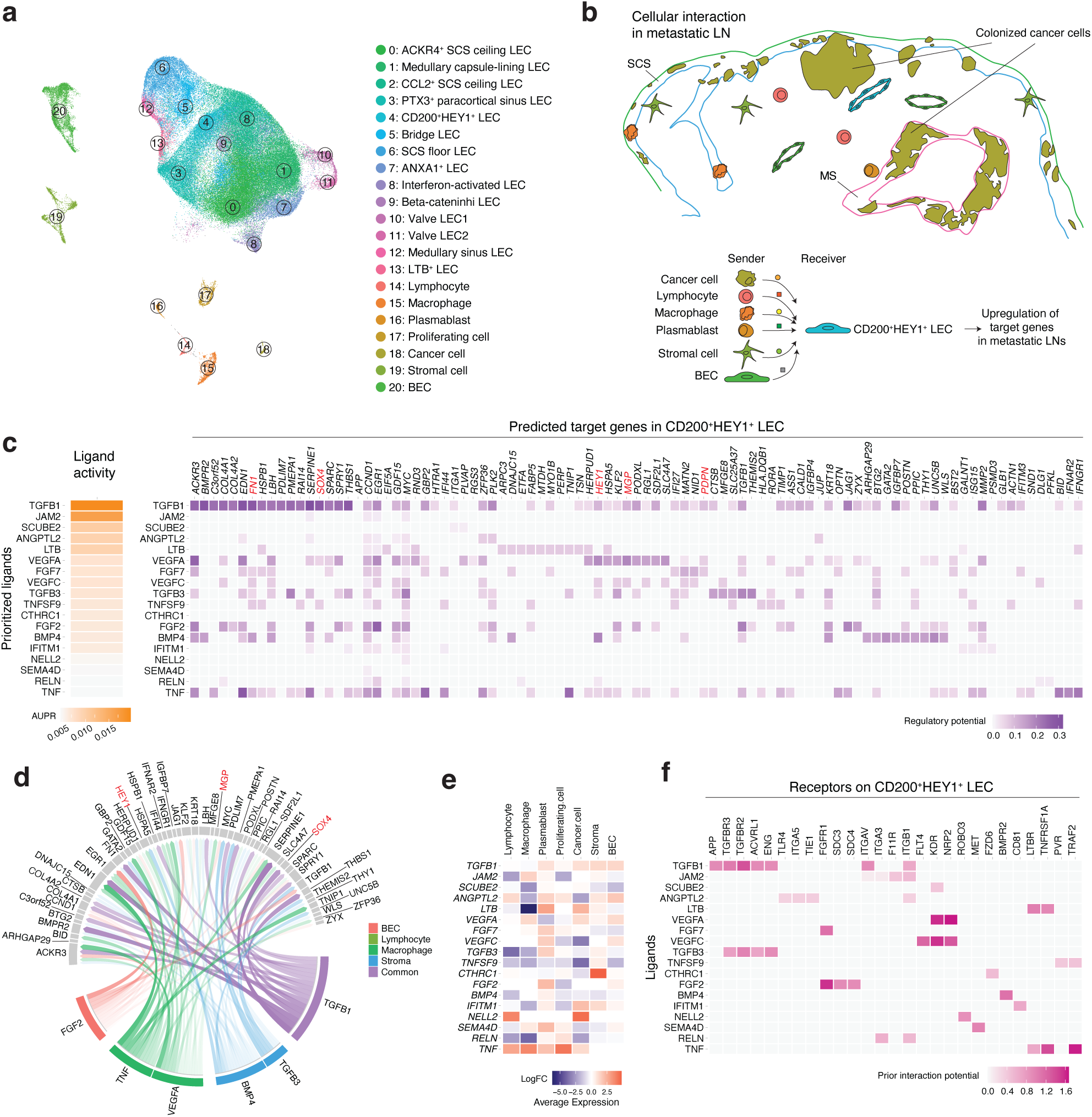
NicheNet predicts the mechanisms driving lymphatic remodeling. **a,** UMAP of 21 cell subsets, including LEC subsets and other cell types in LNs. **b,** A schematic representation of the NicheNet analysis of intercellular communications that induce target genes in CD200^+^ HEY1^+^ LECs in metastatic LNs. CD200^+^ HEY1^+^ LECs were set as the receiver, and the other cell types in LNs were set as the sender. **c,** Predicted top ligands and their target genes in the CD200^+^ HEY1^+^ LECs. AUPR: area under the precision-recall curve. **d,** Circos plot showing the links between predicted ligands from LN cells and their potential target genes in CD200^+^ HEY1^+^ LECs. Low weight links were removed for clarity. **e,** Relative expression of predicted targets in metastatic or distant LNs. **f,** Potential receptors expressed by CD200^+^ HEY1^+^ LECs associated with each predicted ligand.

### Breast Cancer Cell Conditioned Media Alter Human LEC Transcriptomes

To determine whether tumor-induced changes in LECs could be mimicked in an *in vitro* system that allows us to further study the observed changes, we used primary human lymphatic endothelial cells (HLECs), which are isolated from human LNs. In particular, our interest was in MGP that was upregulated in metastatic LN lymphatics of all patients and whose function on lymphatics has remained unexplored. The HLECs were exposed to conditioned culture media (CM) from three different breast cancer cell lines, MCF-7, T47D, and MDA-MB231 (Figure 5a). Both MCF-7 and T47D are estrogen and progesterone positive luminal subtypes of breast cancer and MDA-MB-231 represents triple-negative breast cancer cells. RNA-seq of tumor-conditioned HLECs revealed changes in the expression of multiple genes, with a similar number of genes being upregulated (101 genes) and downregulated (99 genes). While some genes were uniquely changed following exposure to CM from only one breast cancer cell line, other genes showed the same changes with several or all CM, as shown in the Venn diagrams in Figure 5b. Although the total number of upregulated and downregulated genes among all three cancer cell lines were comparable, MDA-MB-231 cells induced the most unique alterations in gene expression. In addition, MDA-MB-231 cells upregulated more genes than they downregulated, whereas this was the opposite with the CM of T47D cells.

**Figure 5.**
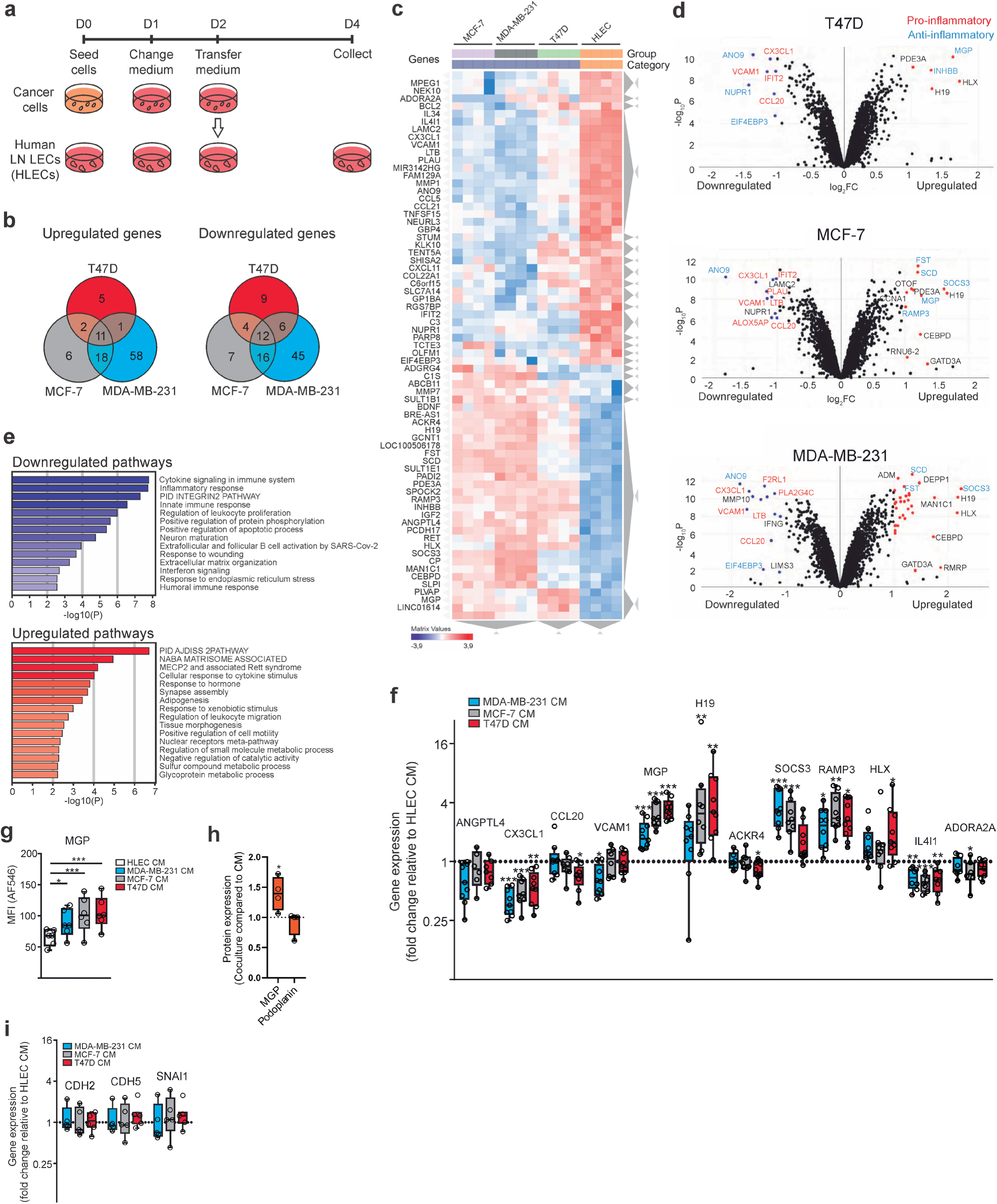
RNA-seq of human LECs exposed to breast cancer cell CM. **a,** Experimental setup for the *in vitro* experiments. Cells were seeded at Day 0 (D0), CM was generated on D1 to D2, and receiving cells were exposed on D2 to D4. **b,** Venn diagrams showing the number of upregulated and downregulated genes obtained by RNA-seq of CM-exposed LECs. **c,** Heatmap showing the top DEGs in the CM-exposed LECs. **d,** Volcano plots depicting genes upregulated and downregulated in LECs after CM exposure. Genes indicated in red are considered mostly pro-inflammatory, whereas genes in blue are anti-inflammatory. **e,** Pathway analysis showing the most significantly downregulated and upregulated pathways (obtained with Metascape and analyzed by the integrated Fisher’s exact test). **f,** Gene expression changes in CM-exposed LECs are shown as determined by qPCR. Data are depicted as Tukey box plots (n = 6–9). Data were analyzed with a linear mixed model and statistical significance adjusted with Dunnett’s multiplicity correction **g,** Protein expression levels of MGP in LECs following CM exposure were determined by flow cytometry, depicted as Tukey box plots (n = 6). **h,** Protein expression of MGP and podoplanin on LECs after coculture of T47D cells with LECs compared to culture with CM. Data were analyzed with 2-way ANOVA and Sidak’s multiple comparison test and are depicted as Tukey box plots. **i,** Expression of *CDH2*, *CDH5*, and *SNAI1* was determined by qPCR in CM-exposed LECs. Data are shown as box plots (n = 5–7). Data were analyzed using one-way ANOVA and statistical significance was adjusted with Dunnett multiplicity correction. The center line of the box plots represents the median, the box represents the 25^th^ to 75^th^ percentiles and the whiskers indicate the inner fences. (*p < 0.05, **p < 0.01, and ***p < 0.001).

To further evaluate these differential gene expression profiles, we implemented unsupervised clustering of genes that were significantly altered in at least two of the three experimental settings based on the RNA-seq transcriptome data (Figure 5c). This resulted in clustering according to the CM used, while also revealing a similar induced profile caused by MCF-7 and MDA-MB-231 CM. Volcano plots for each CM are shown in Figure 5d, where the most altered genes are indicated. They include *MGP*, *SOCS3*, *H19*, and *CEBPD* (upregulated) and *ANO9*, *CX3CL1*, and *VCAM1* (downregulated). In addition, genes associated with a general pro-inflammatory phenotype (shown in red) were downregulated in HLECs cultured with cancer cell CM, whereas anti-inflammatory genes (shown in blue) were upregulated, indicating that soluble factors derived from cancer cells may induce a transition in LECs from a pro-inflammatory to an anti-inflammatory state. To better understand the observed changes in gene expression, we performed a pathway analysis (Figure 5e). Most significantly, pathways involved with inflammation, such as “cytokine signaling in immune system”, “inflammatory response”, or “innate immune response” were downregulated. In contrast, the most upregulated pathways include pathways involved in cell adherence and stability, the matrisome, or cellular response to different stimuli (e.g., “PID AJDISS 2PATHWAY (Posttranslational regulation of adherence junction stability and disassembly)” or “cellular response to cytokine stimulus”).

To verify the observed changes in RNA-seq, we implemented qPCR and flow cytometry assays for selected hits (Figure 5f and g). This confirmed the overall pattern of expression changes, and among them, MGP, receptor activity modifying protein 3 (RAMP3), and interleukin 4 inducible 1 (IL4I1) showed the most consistent alterations. The changes in MGP were the most significant with CM from T47D cells and the least with the CM from MDA-MB-231 cells. We further assessed, using T47D cells, whether direct co-cultures cause similar or stronger phenotypic changes than CM. Indeed, direct co-culture increased MGP levels significantly more than CM, whereas comparable changes were not observed for PDPN (Figure 5h).

MGP was, as in our single-cell-sequencing data, again one of the molecules with the most significant changes. Despite existing beliefs that tumors induce LEC proliferation in murine models, we observed no differences in cell proliferation upon incubation with various media (Supplemental Figure 8).

Blood vessel endothelial cells transform into mesenchymal cells in certain pathologies such as cardiac fibrosis, atherosclerosis, vascular calcification, and cancer. This process is known as the endothelial-to-mesenchymal transition (EndMT) and may play a role in angiogenesis, generation of cancer-associated fibroblasts, and cancer metastasis ^40^. We sought to determine whether this process was induced in LECs by exposure to the CM. As seen in Fig. 5i, this was not the case as none of the EndMT indicative genes *CDH2*, *CDH5*, and *SNAI1* were altered in HLECs following CM exposure.

### Tumor-Induced Transcriptional Changes in LECs are Mediated in Part by VEGFs and TGF-β

Next, we searched for cancer cell-derived factors that can induce transcriptional changes in LECs. The prime candidate cytokines for testing were selected from the published list of cytokines secreted by the breast cancer cell lines used in our study ^41,42^. The selected candidate cytokines were those secreted by all the lines. To verify their effect, we tested whether recombinant VEGF165 (a major spliced isoform of VEGF-A), VEGF-C, TGF-β, EGF, GM-CSF, M-CSF, or G-CSF affect MGP expression. TGF-β, VEGF-A, and VEGF-C are among NicheNet-predicted ligands that may induce lymphatic remodeling. Indeed, VEGF-165, VEGF-C, and TGF-β dose-dependently increased MGP (Figure 6A). The other cytokines did not have an effect (Figure 6a; Supplemental Figure 9a).

**Figure 6.**
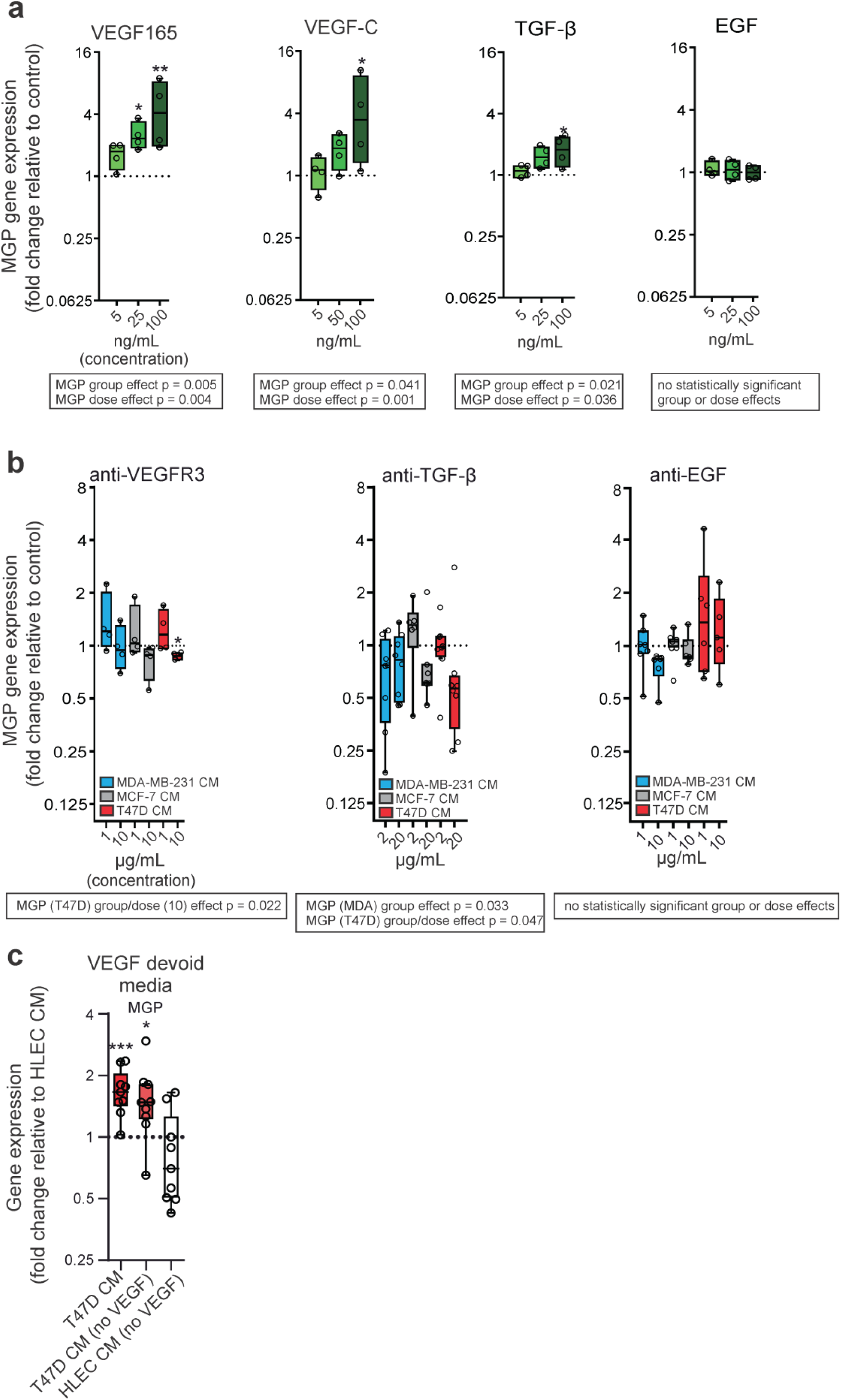
MGP expression in LN LECs is affected by VEGF and TGF-β. **a,** Expression of *MGP* in LECs after direct exposure to recombinant VEGF165 (5, 25, or 100 ng/mL), VEGF-C (5, 50, or 100 ng/mL), TGF-β (5, 25, or 100 ng/mL), or EGF (5, 25, or 100 ng/mL) are shown (n = 3–4). Data are depicted as Tukey box plots and analyzed using linear mixed models fitted separately for each parameter with group (recombinant vs control) and dose and their interaction as fixed effects. **b,** Gene expression changes in modified CM-exposed LECs are shown as determined by qPCR. CM was generated in the presence of antibodies against VEGFR3 (1 or 10 µg/mL, n = 3–4), TGF-β (2 or 20 µg/mL, n = 5–10), and EGF (1 or 10 µg/mL, n = 5–7), compared to Isotype control exposed samples and data are depicted as Tukey box plots showing relative gene changes and analyzed using linear mixed models fitted separately for each parameter with group (antibody vs control) and dose and their interaction as fixed effects. **c,** *MGP* expression determined by qPCR. CM generated with culture media devoid of VEGF supplement was used. Data are shown as Tukey box plots (n = 10). Data were analyzed using a linear mixed model. The center line of the box plots represents the median, the box the 25^th^ to 75^th^ percentiles and the whiskers the inner fences. (*p < 0.05, **p < 0.01, and ***p < 0.001). Statistics of group and dose effects are presented within the boxes; differences in comparison to the controls (defined as 1) are indicated by stars.

We also added blocking antibodies against VEGFR3, TGF-β, EGF, PDGF, and adrenomedullin to cancer cells before starting CM generation, which was subsequently transferred to the HLECs. The addition of blocking antibodies against VEGFR3 and TGF-β, led to reduced induction (or reduction) of MGP in the presence of MDA and T47D CM (Figure 6b). In contrast, we observed no statistically significant changes when blocking antibodies targeting EGF or PDGF were used (Figure 6b; Supplemental Figure 9b). Similarly, the PLA2 inhibitor AACOCF3 did not alter MGP expression. Instead, the STAT1 inhibitor fludarabine dose-dependently decreased MGP in T47D CM, and inhibition of adrenomedullin increased MGP in MDA CM (Supplemental Figure 9b). We further confirmed that cancer cell-secreted VEGFs are responsible for the effect and not VEGF-containing culture medium by using culture medium without VEGFs (Figure 6c). These observations are in line with NicheNet analysis results of scRNA-seq data (Figure 4), and indicate that the cytokine milieu in metastatic LNs could play a crucial role in shaping the transcriptional programs of LECs.

We then focused on MGP, because it was upregulated in the sentinel node LECs of all patients and on the HLECs by CM of all breast cancer cell lines *in vitro*. Moreover, its expression has been associated to the outcome of several cancers ^43^, but its functional role in lymphatics has remained unexplored. We tested whether MGP could regulate its own expression levels. Therefore, we used an anti-MGP antibody while generating CM from T47D cells, before exposing HLECs to it. As seen in Fig. 7a, this did not have a noticeable effect because MGP expression levels were still elevated. Similarly, also exposing HLECs directly to an anti-MGP antibody or a recombinant form of the protein did not alter its mRNA expression (Figure 7b).

**Figure 7.**
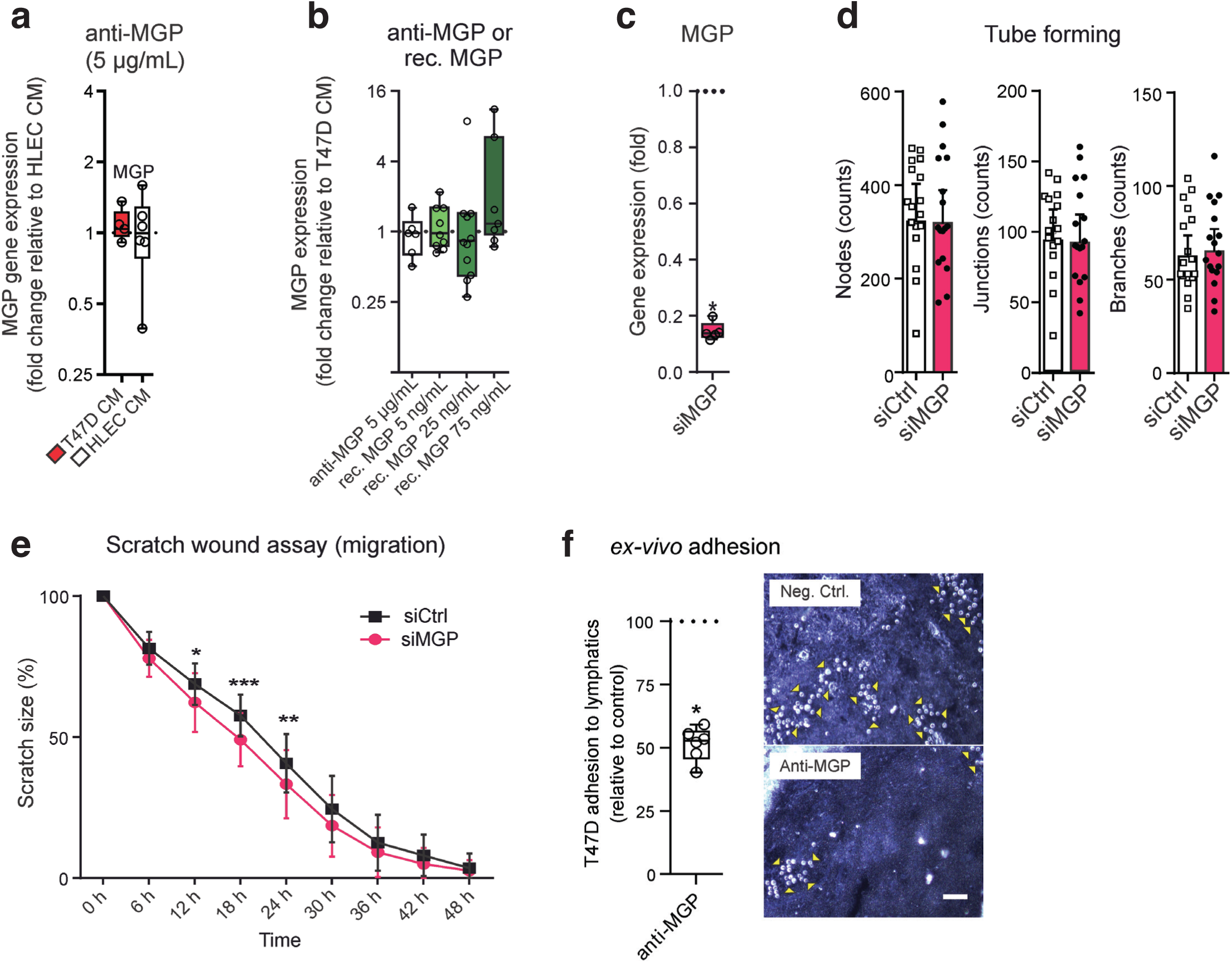
MGP contributes to breast cancer cell adhesion. **a,** Expression of *MGP* in LECs following CM treatment with anti-MGP antibody. Data are shown as Tukey box plots (n = 5– 7). **b,** *MGP* expression in LECs after exposure to anti-MGP antibody (5 µg/mL) or recombinant MGP (5, 25, or 75 ng/mL). Data are shown as Tukey box plots (n = 6–11). **c,** Expression of *MGP* determined by qPCR. Data of siRNA-silenced LECs are shown as Tukey box plots (n = 5) and analyzed with Mann–Whitney U-test. **d,** Tube formation was determined by the number of nodes, junctions, and branches between MGP-silenced and control LECs. Data are shown as the geometric mean with 95% CI (n = 17). **e,** Scratch assay of MGP-silenced and control LECs over the course of 2 d. Data are shown as mean with 95% CI (n = 13) and were analyzed using two-way ANOVA and Sidak’s multiple comparison test. **f,** *Ex vivo* adhesion assay measuring binding of T47D cells to lymphatic sinuses of metastatic LNs after treating LN sections with anti-MGP antibody or negative control antibody. Binding after negative control antibody was defined as 100% due to day-to-day variation in absolute numbers. Total of 1445 cells were found binding to lymphatic sinuses in control antibody treated sections, while 767 cells were bound on anti-MGP treated ones. Binding after anti-MGP treatment is shown as Tukey box plot (n=6) and was analyzed using the ratio paired t-test. An example of T47D cell binding after negative control antibody and anti-MGP treatments to the same area of a metastatic LN is shown below. As the adherent cells (visible as white round circles, some pointed out by yellow arrowheads) are laying on the top of the tissue section, the focus layer of the photograph was chosen to clearly visualize the adherent cells. Scale bar 50 μm. For box plots, the center line represents the median, the box represents the 25^th^ to 75^th^ percentiles and the whiskers indicate the inner fences. (*p < 0.05, **p < 0.01, and ***p < 0.001).

### MGP is Involved in Cancer Cell Adhesion to LECs

Because MGP expression in HLECs was upregulated by cancer conditioning in both the patients’ scRNA-seq data as well as in our *in vitro* assays, we wanted to determine the functions of MGP on HLECs. We used siRNA to silence it and could reduce its expression at the RNA level by approximately 85% (Figure 7c). It has been shown that MGP inhibits EndMT in the blood vasculature by inhibiting bone morphogenetic protein ^44^. Thus, we next tested whether MGP silencing induced EndMT. This was not the case as the observable small changes in gene expression were contraindicative (Supplemental Figure 10). Using the silenced cells, we performed tube-forming assays, which revealed that the tube-forming properties of control and MGP-silenced HLECs were identical (Figure 7d). In contrast, when we performed scratch assays, faster migration of MGP-silenced HLECs was observed (Figure 7e), indicating that MGP inhibits LEC migration.

The impact of MGP on LEC functions was also observed in our cancer cell adhesion experiments. We tested the role of MGP on cancer cell adhesion to LN lymphatics using metastatic lymph node sections of four patients and T47D cells in *ex vivo* adhesion assays. In these assays, the anti-MGP antibody significantly inhibited T47D cell binding to lymphatic sinuses (Figure 7f).

Overall, these data indicate that breast tumor-induced MGP upregulation on LECs may play roles in inhibiting LEC migration, but increases their interaction with breast cancer cells. Thus, TGF-β and VEGF-induced upregulation of MGP in LECs may be a part of the dissemination mechanism of breast cancer cells in patients.

## DISCUSSION

In this study, we observed significant alterations in LN lymphatics draining breast cancer using scRNA-seq analyses. Metastatic LNs showed a loss of specific LEC subsets, such as the SCS floor and medullary sinus LECs. However, they displayed the emergence of certain abnormal LEC subsets, including CD200^+^ HEY1^+^ LECs. This LEC remodeling signature was consistently observed in multiple patients. Among the upregulated molecules, MGP expression was notably high in metastatic LN LECs of patients with breast cancer, a characteristic that was also observed in cultured LN LECs in the presence of CM of breast cancer cells. MGP silencing in LECs increased LEC migration and reduced cancer cell adhesion. These findings demonstrate that lymphatic remodeling in sentinel LNs is intimately linked to cancer progression.

In previous studies, we reported the heterogeneity of LECs in human LNs, identifying six distinct types of LN LECs ^15^. In the present study, we discovered additional subsets (total 14 clusters) in non-metastatic LNs. This disparity may stem from several factors. Notably, our current report incorporates the analysis of a larger number of LECs, uses a newer, improved version of 10x Genomics kits and Cell Ranger version 3 instead of v2. We also used updated genome reference GRCh38, in contrast to GRCh37 used in the previous study. This may have helped to align the sequenced read more accurately. For example, PECAM1 was not detected in the previous dataset, but was successfully detected in the current dataset. We currently also employ a higher number of principal component analysis (PCA) dimensions to identify transcriptionally distinct subsets compared with our previous analysis. Moreover, we utilized the more recently developed Seurat version 4 integration tools, which surpass the capabilities of the canonical correlation analysis (CCA)-based data integration tools used in our prior study. These new tools allowed us to apply a higher number of PCA dimensions while effectively mitigating technical batch effects between the samples. Additionally, since the clustering parameters were selected manually to identify 14 LEC clusters, there may be additional cellular subsets or states that have not been captured.

In metastatic LNs, there was an observed increase in capillary-like CD200^+^ HEY1^+^ LECs and paracortical sinus, whereas SCS floor and medullary sinus LECs were reduced. Biglycan, encoded by *BGN*, is a small leucine-rich repeat proteoglycan and a constituent of the extracellular matrix. Notably, biglycan was discovered to be significantly more abundant in the blood ECs of tumors than in ECs in normal tissues ^45^. Biglycan is expressed in several cancers, and its expression on fibroblasts and cancer cells is induced by TGF-β ^46^. Our analysis showed this subset also expresses the immunosuppressive molecule CD200. CD200 is expressed by multiple cell types such as epithelial cells, BECs and fibroblasts, and delivers an inhibitory signal in the inflammation by binding receptors, which are found in myeloid lineage cells and innate lymphoid cells ^47–50^. The expression of CD200 in LECs may provide an inhibitory signal to macrophages or dendritic cells, which are often found in lymphatic sinuses, to evade tumor immunity. Another increased LEC subset in metastatic LNs is paracortical sinus LEC and this subset exhibits a markedly higher expression of VEGFR3 compared with other LEC subsets in humans, potentially facilitating faster growth in response to VEGF-C produced by the tumor. Given that these paracortical sinuses are the exit points for lymphocytes in LNs and that we detected individual cancer cells within CD200^+^ lymphatics, the robust enrichment of these subsets in metastatic LNs may be indicative of the efficiency of tumor egress through the LNs. The reduced subsets of SCS floor and medullary sinus LECs exhibit high expression of molecules involved in immunological defenses and inflammation compared with other LN LEC subsets ^15^. These sinus LECs produce CSF1 and are crucial for maintaining SCS and medullary sinus macrophages ^34^, elements vital for bacteria ^51^ and virus infection ^52^, as well as tumor immunity ^35^. Hence, our study indicates that tumor metastasis within LNs instigates a transition from immunologically active LEC subsets such as SCS floor LECs to those that are immunosuppressive and conductive to tumor metastasis.

Lymphatics serve not only as conduits for immune and cancer cell migration but are also active participants in the immune response through antigen presentation. It has been demonstrated that LN LECs express self-antigens and present them on MHC class I and II molecules ^53^. Despite the lack of costimulatory molecules, LN LECs do exhibit high expression of inhibitory molecules such as PD-L1, thereby inducing peripheral tolerance by prompting apoptosis in cytotoxic memory CD8^+^ T cells ^54^ and driving PD-1 expression of CD8^+^ T cells ^55^. Our prior research showed that, in particular, SCS floor and medullary sinus LECs strongly express PD-L1 in both mice and humans ^15,16^. Given the selective expression of PD-L1 on LECs, but not other cell types in LNs, the reduction of PD-L1^+^ SCS floor LEC subsets and the accumulation of PD-L1^−^ LEC subsets in metastatic LNs may significant impact the efficacy of immune checkpoint therapy, such as anti-PD-L1 agents like Atezolizumanb and Durvalumab, cancer patients.

Based on the current patient cohort and data, it is impossible to predict, which of the found changes have an impact on clinical outcome. This is due to the relatively small number of patients without sufficiently long follow-up times. However, a recent study reported immunohistochemical analyses of LNs of breast cancer patients and found that high PDPN expression in the lymphatics of sentinel LNs (also found by us, Fig. 3) is a prognostic parameter for worse overall survival, and independent of tumor size, nodal status and age at the surgery ^56^.

Similar to the results obtained from single-cell sequencing of patient samples, we found gene alterations caused by cancer cell CM in *in vitro* experiments. As we aimed to find universal effects on gene regulation, we focused on changes shared between the different cancer cell lines. Nevertheless, it was interesting to notice that MDA-MB-231 cells, the cell line representing triple-negative breast cancer, caused the most significant gene alterations. This agrees with the clinical behavior of triple-negative breast cancer as it has a more aggressive and disruptive phenotype compared with luminal cancer types (such as T47D and MCF-7) ^57^. When looking at common changes induced by CM, we observed a shift of the endothelial cells to a more anti-inflammatory state, a finding that has previously been found in other cell types such as macrophages and T cells ^58–60^. This phenotypical change is therefore likely beneficial for tumor growth and expansion.

MGP was constantly upregulated both in the lymphatics of patients with cancer and *in vitro* LEC cultures. This is in line with a recent study analyzing changes in co-cultures of human melanoma cell lines and LECs using scRNA seq ^61^. In those analyses, *MGP* together with *BGN, MGP, SOX4* and *MFAP2*, which are also among our best hits were significantly upregulated. Although the literature is mostly focused on the role of MGP in calcification and vascular morphogenesis ^62,63^, it has also been associated with the outcome of various cancers. However, the role of MGP in terms of cancer appears complicated, as not only can its expression be considered both beneficial and detrimental, depending on the cancer type, but also its molecular effect seems to vary ^64–68^. Furthermore, MGP expression correlates differently to metastatic spread depending on the tumor type and high expression associates with the metastatic spread in breast cancer ^43^.

As the role of MGP in LECs is not known, we wanted to focus on its functional importance especially on LECs. Silencing of MGP increased migration in the scratch assay, suggesting a migration inhibitory function of MGP. In addition, cancer cell adhesion was decreased in MGP-inhibited LECs. Such a function could therefore be an indication of a potentially induced facilitation of cancer cell spreading. In fact, Zandueta et al. have reported MGP on osteosarcoma cells to promote their adhesion to and transmigration via endothelial cells leading to enhanced metastasis ^69^. This together with our results concerning LECs suggests that MGP can mediate adhesion in different cell types. Although MGP is a secreted K vitamin dependent protein, it may use its GLA-domain to remain bound to the cell membrane as GLA-proteins have been shown to do^70^ Comparable to what we have seen in our study regarding lymphatics, MGP has been described as a molecule that suppresses angiogenic sprouting as well as angiogenesis ^71^, while others have reported conflicting results describing it as an angiogenesis-promoting molecule ^72^. It therefore seems to become clear that the function and effects of MGP are dependent on its environment, the cell type, as well as the condition where it is studied. Hence, MGP exerts diverse roles with regards to the tumor environment and tumor type and more in-depth research is required to pinpoint its specific role and importance.

When exploring the potential factors enhancing MGP expression, we first chose candidates among the reported factors secreted by cancer cell lines and found that VEGF165, VEGF-C, and TGF-β upregulate MGP expression in LECs. We also witnessed reduced expression of MGP after using antibodies against VEGFR3 in the CM of T47D cells, and a trend with CM from other cancer cells. Similarly, we found reduced MGP with anti-TGF-β in the CM of MDA and T47D cells. These findings are in line with earlier observations, in which VEGF upregulated MGP on retinal endothelial cells ^73^ and TGF-β on bovine aortic endothelial cells ^74^. Results of some previous work not utilizing LECs indicated that MGP expression is upregulated on bone marrow endothelial progenitor cells by PDGF ^75^, upregulated by TGF-β in embryonic mouse lungs ^76^, and downregulated in the trabecular meshwork by TGF-β ^77^. These findings indicate that a complex regulation pattern for MGP exists and is dependent on the cell type, their density in cell cultures, and treatment regimens ^78^. Complexity is further supported by our present observation that CMs reduced gene expression of *IL4I1*, while with recombinant VEGF165 and VEGF-C we detected an increase. This may be due to multiple factors in the CM causing different effects. Although we studied the importance of different factors *in vitro*, it is likely that these or similar cytokines are also responsible for the observed changes in patients *in vivo*.

There are limitations in this study. We cannot completely exclude the possibility that the non-draining LNs would be exposed to some tumor-derived soluble factors, which originate from the draining node being in the same chain. Because there are 20 or even more axillary nodes, it seems unlikely that marked amounts of cancer-secreted factors would end up to the distal node. In addition, ethical requirements do not allow more invasive experimental approaches including the determination of the time course leading to the alterations in the sentinel nodes. Thus, we believe that the changes in the draining LNs are caused by cancer derived factors, such as those carried by extracellular vesicles via lymphatics and described in the literature, ^79^ and/or the presence of cancer cells via cell-cell contacts.

Another potential limitation of our study is, that once LN LECs are isolated and cultured, they dedifferentiate and no longer accurately represent individual clusters *in vivo*. However, they can still serve as a tool to analyze the induction and function of various genes or proteins, as we were able to identify several cancer-induced changes that were shared between the HLECs and certain *in vivo* clusters. In this context we analyzed the function of MGP, which was highly upregulated both in lymphatics *in vivo* and HLECs *in vitro* and were able to discover its adhesive function supporting cancer cell binding to LECs.

In summary, our study demonstrates that LEC cell heterogeneity in LNs is much greater than previously reported. Moreover, we show that breast cancer modifies the lymphatics in sentinel LNs with respect to their phenotype and function. MGP appeared as the top molecule in LECs upregulated by both different types of breast cancers *in vivo* and cancer cell lines *in vitro*. Based on the functional studies, we propose that cancer-promoting MGP expression in LECs has the potential to contribute to cancer behavior.

## Acknowledgements

We thank Teppo Huttunen from Estimates for statistical analyses, Sari Mäki and Riikka Sjöroos for technical help, Joe Hettinger for revising the language, Bishwa Ghimire for his advice on bioinformatic analyses, and Prof. Kari Alitalo for providing us with anti-VEGFR3 antibody. This work was financed by the Research Council of Finland (earlier the Finnish Academy), the Finnish Cancer Foundation, Sigrid Juselius Foundation, Jane and Aatos Erkko Foundation and Sakari Alhopuro Foundation.

## METHODS

### Study design

This study was designed to analyze how tumor can modify the lymphatics in the sentinel lymph nodes. We used a unique patient material by comparing the lymphatics in the sentinel metastatic lymph nodes to distant nonmetastatic lymph nodes not draining the tumor area in the same patients and thus, excluded the patient specific differences. Single cell sequencing combined with immunohistochemistry revealed significant changes in lymphatic endothelial cells caused by the cancer. These changes were then studied in a more simplified settings using breast cancer cell lines and lymphatic endothelial cells in culture conditions. The functional properties and inducing factors of the best hit molecule upregulated by cancer both *in vivo* and *in vitro* were studied using tube forming, scratch wound (migration) and adhesion assays.

### Human Samples

Axillary LNs were collected from patients with breast cancer undergoing mastectomy and axillary LN removal at Turku University Hospital. Sentinel LNs as well as non-metastatic, distant LNs that do not drain the tumor were obtained. The draining lymph nodes were identified using preoperative lymphoscintigraphy with 99mTC nano-colloid and perioperative Patent Blue that is injected into the tumor helping to visualize the lymphatics and the draining node. The draining nodes, typically 1-3 are then detected based on the blue color and the radioactive signal (detected by a hand-held gamma counter). The presence of tumor cells in metastatic and distant LNs was determined by pathological assessment and flow cytometry analysis of pan-cytokeratin staining before single-cell sequencing. The collection was conducted under the license EMTK: 132/2016. Written informed consent was obtained from each tissue donor. The samples were anonymized and handled according to the ethical guidelines established by the University of Turku. The samples were used with the permission of the Institutional Review Board of Medicolegal Affairs (Helsinki and Turku, Finland).

### LN LEC Isolation

scRNA-seq of LN LECs was conducted following previously described methods (13). Immediately after surgery, human LNs were placed in RPMI, cut into small pieces, and digested for 1 h with RPMI containing 0.2 mg/mL collagenase P, 0.8 mg/mL dispase, and 0.1 mg/mL DNase. To confirm the presence of cancer cells in the LNs, a portion of the single-cell suspension was stained with AF488 anti-pan-cytokeratin antibody (ThermoFisher Scientific, MA5-18156) after permeabilization. Pan-cytokeratin–positive or –negative LNs were considered as metastatic or distant LNs, respectively. CD45^+^ cells were depleted using the EasySep Human CD45 Depletion Kit (Stem Cell Technologies), and the single-cell suspension was incubated with PE anti-PDPN (Biolegend, 337004), AF488 anti-CD45 (Biolegend, 304019), APC anti-CD31 (Biolegend, 303115), and LIVE/DEAD Fixable Near-IR Dead Cell Stain Kit (ThermoFisher Scientific, L10119) for 30 min. Live CD45^−^ PDPN^+^ CD31^+^ LECs were sorted using SH800S (Sony) in DMEM medium containing 10% fetal calf serum (FCS).

### scRNA-seq and Data Preprocessing

Sorted LECs were promptly counted manually and processed following 10x Genomics guidelines. scRNA-seq libraries were prepared according to the manufacturer’s instructions using the Chromium Single Cell 3′ Library and Gel Bead Kit v2 and v3 (10x Genomics, 120237) along with the Chromium Single Cell A Chip Kit. In brief, cells were mixed with reverse transcriptase master mix and partitioned into nanoliter-scale gel bead-in-emulsions (GEMs) using 10x Genomics GemCode Technology. Barcoded cDNA was purified and amplified by PCR. P5, P7, a sample index (i7, 10x Genomics, 120262), and R2 (read 2 primer sequence) were added during library preparation. Sequencing was performed using the Illumina HiSeq3000 or NovaSeq platforms. Each sample was sequenced to achieve an average depth of 50,000 reads per cell. Postprocessing, including demultiplexing, read alignment, and quality control, was performed at the Medical Bioinformatics Centre at the University of Turku using the 10x Genomics Cell Ranger package (version 3.0.1 or 3.1.0). The data are accessible at the Gene Expression Omnibus (GEO) repository (#GSE248214).

### scRNA-seq Data Processing and Clustering

Preprocessed data were analyzed using Seurat (version 4.3) for graph-based clustering and differential gene expression analysis. For quality control, we filtered out cells with unique feature counts over 6,000 or under 200, and cells with more than 12.5% mitochondrial counts. For integration, each dataset was normalized and 2,000 variable genes in each dataset were identified using the “NormalizeData” and “FindVariableFeatures” functions, respectively. Shared highly variable genes across datasets were identified using the “SelectIntegrationFeatures” function. Integration anchors were identified on the basis of these genes using CCA ^22^ as implemented in the “FindIntegrationAnchors” function. The data were then integrated using “IntegrateData()” and scaled using “ScaleData”. PCA and uniform manifold approximation and projection (UMAP) dimension reduction with 30 principal components were performed. A nearest-neighbor graph using the 30 dimensions of the PCA reduction was calculated using “FindNeighbors”, followed by clustering using “FindClusters” with a resolution of 0.5. The clustering was visualized with UMAP using “DimPlot”. A cluster expressing heat-shock proteins such as HSPA1A/B was removed from further analysis, as these gene were shown to be affected by cell dissociation ^23^. Another cluster with high expression of mitochondrial genes like *MT-CO3* and *MT-CO1* was also excluded from further analysis. Markers used to phenotype cells in human LNs included PROX1 and FLT4 (LECs), JAM2 (BECs), COL1A1 (stromal cells), PTPRC (leukocytes), KRT19 (cancer cells), and MKI67 (proliferating cells). The PROX1^+^ LEC subsets was gated and subclustered by re-identifying integration anchors within datasets as described above.

### Differential Expression Analysis Using scRNA-seq Data

Differentially expression genes (DEGs) between clusters, metastatic state, and patients were identified using “FindAllMarkers”. The “Min.pct” parameter (minimum percentage of the gene-expressing cells in each cluster) was set to 0.25 (25%), and “thresh.use” (minimum fold change in the gene expression between each cluster and all other clusters) was set to 0.25 (log2FC). Violin plots and heatmaps were generated using Seurat’s “VlnPlot” and “DoHeatmap” functions or Scillus’s “plot_heatmap” function. Pathway enrichment analysis of DEGs was performed using Enricher ^24^.

### Differential Abundance Testing

The R package Milo (ver.1.8.1) was used for performing differential abundance testing on the k-nearest neighbor (KNN) graph ^25^. The Seurat dataset was converted into a single cell experiment format and x, y coordinates of UMAP were imported from the Seurat dataset. Covariates in the differential abundance testing were ‘original.ident’ (individual sample identity) and ‘status’ (distant or metastatic). Milo analysis was performed using the standard workflow, and the KNN graph was generated from the latent space available from Seurat.

### Trajectory Analysis of LECs

We performed a pseudo time trajectory analysis of LECs using the Monocle3 package ^26,27^. Monocle3 projects cells onto a low-dimensional space using UMAP, groups similar cells together using the Louvain algorithm, and then merges adjacent groups and resolves the paths for individual cells ^27^. The setting to include a circular path was turned off. Consistent with an earlier study ^15^, we selected the root node from within ACKR4^+^ SCS ceiling LECs.

### Prediction of Cell Interactions

NicheNet method was used for analyses of cell-cell communication ^28^. NicheNet predicts which ligands from sender cells regulate the expression of target genes in receiver cells by integrating scRNA-seq data with a prior knowledge of signaling and gene regulatory networks. In this study, we initially used NicheNet to identify potential communication between combinations of all 21 cell types exhaustively. We then performed a separate analysis, in which CD200^+^ HEY1^+^ LECs and SCS floor LECs were defined as receivers and lymphocytes, macrophages, plasma blasts, proliferating cells, cancer cells, stromal cells and BECs were defined as senders. For CD200^+^ HEY1^+^ LECs, we considered upregulated genes and for SCS floor LECs downregulated genes as genes of interest. In both analyses, we used the following cut-offs: log fold change of 0.25, fraction of cells expressing the gene of 0.1, maximum number of ligands of 20, maximum number of targets of 100 and ligand-target score of 0.33.

### Immunohistochemical Staining

Fresh human LNs were embedded in OCT compound (Sakura) and frozen on dry ice. The LNs were sectioned at a thickness of 6 μm using a cryostat and fixed with acetone at −20°C for 5 min. The sections were incubated with 10% FCS added to 0.5% BSA in PBS at room temperature for 1 h to prevent nonspecific binding. Then, they were covered with primary antibodies diluted in the same buffer (10% FCS + 0.5% BSA in PBS) and left for overnight incubation at +4°C. The following primary antibodies were used: goat anti-human PROX1 (R&D Systems AF2727), rabbit anti-human MARCO (Atlas Antibodies, HPA063793), AF488 mouse anti-pan-cytokeratin (Thermo Fisher Scientific, MA5-18156), mouse anti-human MGP (Novus Biologicals, NBP2-45844), and BV421 mouse anti-human CD200 (Biolegend, 329209). Subsequently, the slides were washed and incubated with the following secondary antibodies for 1 h at room temperature: AF488 donkey anti-goat IgG (Thermo Fisher Scientific, A32814), AF647 donkey anti-goat IgG (Thermo Fisher Scientific, A21447), AF555 donkey anti-rabbit IgG (Thermo Fisher Scientific, A32794), and AF647 donkey anti-rabbit IgG (Thermo Fisher Scientific, A32795). After washing with PBS, the sections were mounted with ProLong Gold Antifade Mounting medium with DAPI (ThermoFisher Scientific, P36931). Images were captured using an LSM880 confocal microscope (Zeiss) or Stellaris 8 Falcon FLIM microscope (Leica), and analyzed and quantified using ImageJ.

### Cell Lines

Human LECs from LNs were obtained from ScienCell (#2500) and CellBiologics (#H-6092) or extracted from patient material as described above and were cultured in their respective media (#1001 and H1168, respectively). MDA-MB-231, T47D, and MCF-7 cells were purchased from ATCC and maintained in the laboratory. T47D, and MCF-7 cells were cultured in DMEM supplemented with 10% FCS, 100 U/mL penicillin, and 100 µg/mL streptomycin. MDA-MB-231 cells were also cultured in the described DMEM media, but additionally had 4 mM L-glutamine and MEM non-essential amino acids (Thermo Fisher, 11140050).

### Generation of Conditioned Media and Cell Conditioning

Cancer cells or HLECs were plated in their respective media and let to adhere for 1d before all media was changed to HLEC media. After 1d, the media was centrifuged at 450g for 10 minutes before adding it to cultured HLECs for 2d. In 12-well plates, 100000 cells were plated for generating, 45000 cells were plated for receiving the conditioned media. In 24-well plates these number were 50000 and 23000, respectively.

### Cell Co-culture of HLECs and T47Ds

HLECs and T47Ds were plated in their culture medium on d0 in small culture flasks or to a 10 cm culture dish respectively. On d1, medium from T47D cells was collected, centrifuged 450g for 10 minutes and transferred to HLECs. To certain HLECs, 1/3 of their cell number was added as T47D cells (∼340000 cells). On d3 the cells were permeabilized, stained for CD31, podoplanin and MGP and analyzed with flow cytometry.

### Silencing MGP

siRNA silencing was performed with the Lipofectamine RNAiMAX reagent (#13778075) and Silencer Select siRNA (s8753) or the negative control #1 following the manufacturer’s instructions but using only 50% of the recommended reagent amounts (ThermoFisher Scientific, Espoo, Finland).

### Tube-Forming Assay

Growth factor-reduced Matrigel (Corning 356231, Espoo, Finland) was plated in a 96-well flat-bottom plate. Once solidified, 10,000 LECs/well were added and cultured for 1 d. For tube-forming assays with CM, cells were exposed to CM for 2 d before performing the assay. CM was also used during the assay duration.

### Adhesion Assays

To study the role of MGP at the protein level in lymph node lymphatics *ex vivo* adhesion assay was performed as described ^29^. Briefly, lymph node sections from four patients were treated with anti-MGP antibody (1:100, Proteintech 10734-1-AP) or a negative control antibody (Rabbit IgG 1:100, Proteintech 30000-0-A) for 30 min. After removing the antibody, the sections were overlaid with T47D cells and incubated in static conditions for 15 min, followed by rotation at 60 rpm for 5 min, and again without rotation for another 15 min at 7°C. The adherent cells were fixed in 1% glutaraldehyde. The number of malignant cells bound to the lymphatic sinusoids was counted under a dark-field illumination microscope (x200; Leitz Aristoplan, Oberkochen, Germany). To be able to compare experiments performed on different days, the results of the inhibition assays are presented as the percentage of binding found in control treated slides, in which the number of cells adherent to the lymphatics in the presence of the control antibody was taken as 100% adherence.

### Reagents used in *In Vitro* Experiments

Reagents were obtained from different manufacturers. VEGF-C (100-20CD), VEGF165 (100-20), MCSF, GM-CSF, and G-CSF were obtained from Peprotech, and MGP (TP760483) was obtained from Origene. AACOCF3 (ab120350) was obtained from Abcam, fludarabine (S1491) from Seleckchem, antibodies targeting adrenomedullin (AF6108), EGF (AB-236-NA), goat IgG control (AB-108-C), PDGF (AB-20-NA), rabbit IgG control (AB-105-C), and VEGF165 (AB-293-NA) from R&D Systems. Anti-MGP (A5439) was obtained from Abclonal and anti-TGF-β (BE0057) and the mouse IgG1 isotype (BE0083) from BioXCell. The anti-VEGFR3 antibody (IMC-3C5) was a kind gift from Kari Alitalo, and the control antibody was obtained from Bio-X-Cell (human IgG1, #BE0297).

### Cancer Cell Proliferation

Proliferation was followed by labeling the cells with 6.5 µM CellTracker Red CMTPX (C34552, Thermo Fisher Scientific) for 40 min and imaged using a Cytation5 (Agilent BioTek) every 7–8 h over a duration of 2 d.

### Flow Cytometric Analysis of MGP

Cells were blocked for 20 min with human Ig on ice, permeabilized with the BD Cytofix/Cytoperm Kit (554714), and sequentially stained for 30 min with the following antibodies: MGP (NBP2-45844) from Novus; negative control, mouse IgG2a (553454) from BD and anti-mouse IgG2a AF546 (A21133) from Invitrogen.

### Scratch Wound Assays

HLECs were plated on fibronectin-coated wells of a 96-well plate (1,800 cells/well), and 1 d later, the cells were siRNA silenced and cultured for 3 d. A scratch was made to confluent wells with a 200 µL pipette tip, and the well was then imaged with an IncuCyte S3 (Sartorius) every 6 h.

### qPCR

RNA was extracted using the NucleoSpin RNA Kit (Macherey-Nagel, Düren, Germany) according to the manufacturer’s instructions. RNA was converted to cDNA using the SuperScript VILO cDNA Synthesis Kit (Thermo Fisher Scientific, Espoo, Finland) before being used in TaqMan Gene Expression Assays (Applied Biosystems, Stockholm, Sweden) with a QuantStudio3 (Applied Biosystems). To evaluate gene expression levels, the 2^(-ddCT)^ method with B2M as a control housekeeping gene was used. The following probes were used: ACKR4 (Hs00664347_s1), ADORA2A (Hs00169123_m1), ANGPTL4 (Hs01101127_m1), B2M (Hs99999907_m1), CCL20 (Hs00355476_m1), CDH2 (Hs00983056_m1), CDH5 (Hs00901465_m1), CX3CL1 (Hs0171086_m1), H19 (Hs00262142_g1), HLX (Hs00172035_m1), IL4I1 (Hs00541746_m1), MGP (Hs00179899_m1), RAMP3 (Hs00389131_m1), SNAI1 (Hs0019559_m1), SOCS3 (Hs02330328_s1), and VCAM1 (Hs01003372_m1).

### RNA-seq

Total RNA was extracted from LECs using the NucleoSpin RNA Kit (Macherey-Nagel), and library preparation was performed according to the TruSeq Stranded mRNA Sample Preparation protocol using the TruSeq Stranded mRNA HT Kit (Illumina). Sequencing was performed using an Illumina NovaSeq 6000 with a 2 50-bp read length at the Finnish Functional Genomics Centre at the University of Turku and Åbo Akademi as well as Biocenter Finland.

### Bioinformatics for RNA-seq

The raw sequencing data were uploaded to the BaseSpace Sequence Hub (Illumina) as fastq files for further analysis. Quality control was performed using the FastQC application of BaseSpace. Subsequently, the sequences were aligned against the human reference genome hg19 (UCSC, RefSeq gene annotation) with the RNA-Seq Alignment application, which uses the STAR aligner for read mapping and salmon for quantification of reference genes and transcripts. Differences in gene expression between the samples were identified with the RNA-Seq Differential Expression application using DESeq2. Genes exhibiting a fold change >2 (log2ratio ≥1 and ≤−1) and q-values (false discovery rate) < 0.05 were selected as DEGs. The RNA-seq data are accessible at GEO under accession number GSE248076.

Further analyses were performed using the web tools from BioJuppies and MetaScape ^30^. In particular, the heatmap was generated using Clustergrammer ^31^, which normalized raw gene counts using the logCPM method, filtered and transformed using the Z-score method.

### Statistical Analyses

Data were analyzed using GraphPad Prism 6.02 or 9 (Graph-Pad Software, San Diego, USA) and SAS version 9.4 (SAS Institute Inc., Cary, NC, USA). The statistical test used is indicated in each figure. Values of *p < 0.05, **p < 0.01, and ***p < 0.001 were considered statistically significant. A linear mixed model for repeated measurements for dose and group effects was used, and when groups were compared at several dose levels, we applied Sidak’s adjustment for multiplicity.

---

**Supplemental table 1. Excel file containing the up and down regulated genes in different clusters.**

