## Supplemental figures 1-10 for "Breast Cancer Remodels Lymphatics in Sentinel Lymph Nodes"

### Supplemental Figure 1

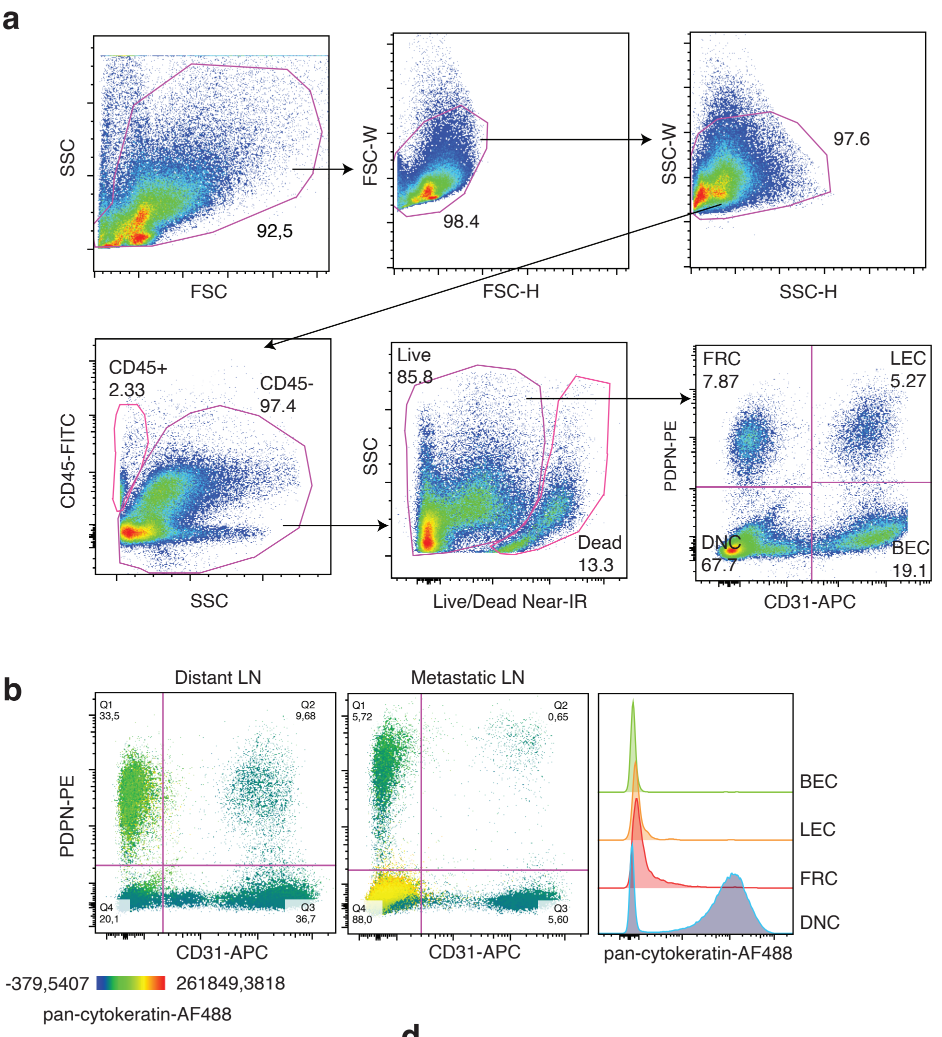


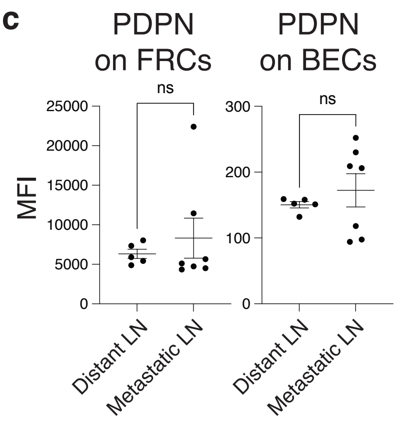


**d**

| Patient ID | LN | LN status | Live cell (%) | FRCs (%) | BECs (%) | LECs (%) |
|---|---|---|---|---|---|---|
| #1 | AXLN1 | Metastatic | 89.3 | 0.96 | 11.3 | 2.23 |
| #1 | AXLN2 | Metastatic | 85.8 | 2.92 | 14 | 5.06 |
| #1 | AXLN3 | Metastatic | 89.7 | 1.01 | 14.8 | 2.5 |
| #1 | AXLN4 | Metastatic | 90.9 | 1.57 | 11.6 | 4.77 |
| #2 | AXLN1 | Metastatic | 68.7 | 17.6 | 15.3 | 2.89 |
| #3 | AXLN1 | Metastatic | 67.5 | 12.9 | 34.6 | 4.41 |
| #3 | AXLN2 | Non-metastatic | 86.9 | 10.9 | 25.9 | 7.26 |
| #4 | AXLN1 | Metastatic | 71.8 | 5.45 | 21 | 5.71 |
| #4 | AXLN2 | Non-metastatic | 90.7 | 9.46 | 38.2 | 5.62 |
| #5 | AXLN1 | Metastatic | 83.4 | 1.86 | 2.26 | 0.33 |
| #5 | AXLN2 | Non-metastatic | 84.2 | 16 | 24.9 | 3.76 |
| #5 | AXLN3 | Non-metastatic | 79.2 | 16.1 | 18.1 | 3.68 |
| #5 | AXLN4 | Metastatic | 79.8 | 7.59 | 2.49 | 0.77 |
| #6 | AXLN1 | Metastatic | 83.6 | 1.94 | 3.82 | 1.95 |
| #6 | AXLN2 | Metastatic | 87.5 | 5.44 | 1.8 | 1.34 |
| #6 | AXLN3 | Non-metastatic | 93.9 | 3.04 | 2.44 | 1.43 |
| #7 | AXLN1 | Metastatic | 87.8 | 24.5 | 22.2 | 2.81 |
| #7 | AXLN2 | Non-metastatic | 94.7 | 16.8 | 44 | 10 |
| #8 | AXLN1 | Metastatic | 93.5 | 9.11 | 21.1 | 4.17 |
| #8 | AXLN2 | Non-metastatic | 87.4 | 15 | 29 | 6 |
| #9 | AXLN1 | Metastatic | 66.5 | 60.9 | 15.3 | 1.21 |
| #9 | AXLN2 | Non-metastatic | 77 | 9.12 | 41.9 | 10.7 |

**Supplemental Figure 1. LEC Isolation. a,** Gating strategy for LEC enrichment. Doublets were excluded, followed by gating of $CD45^-$ cells. Among Live/Dead Near-IR-negative live cells, $PDPN^+$ $CD31^+$ double-positive cells were selected and sorted. **b,** $PDPN^-CD31^-$ double-negative cells (DNCs) include cancer cells. Dot plots (left) display the intensity of pan-cytokeratin expression (indicated by color). Histograms (right) show pan-cytokeratin expression in FRCs ($PDPN^+CD31^-$), LECs ($PDPN^+CD31^+$), BECs ($PDPN^-CD31^+$), and DNCs ($PDPN^-CD31^-$) cells. **c,** PDPN expression in FRCs and BECs from distant and metastatic LNs (mean±SEM, two-tailed, unpaired Student's t-test). **d,** Frequencies of live cells within the $CD45^-$ fraction and the proportions of FRCs, BECs, and LECs among live cells across all samples used in this scRNA-seq study.

#### Supplemental Figure 2

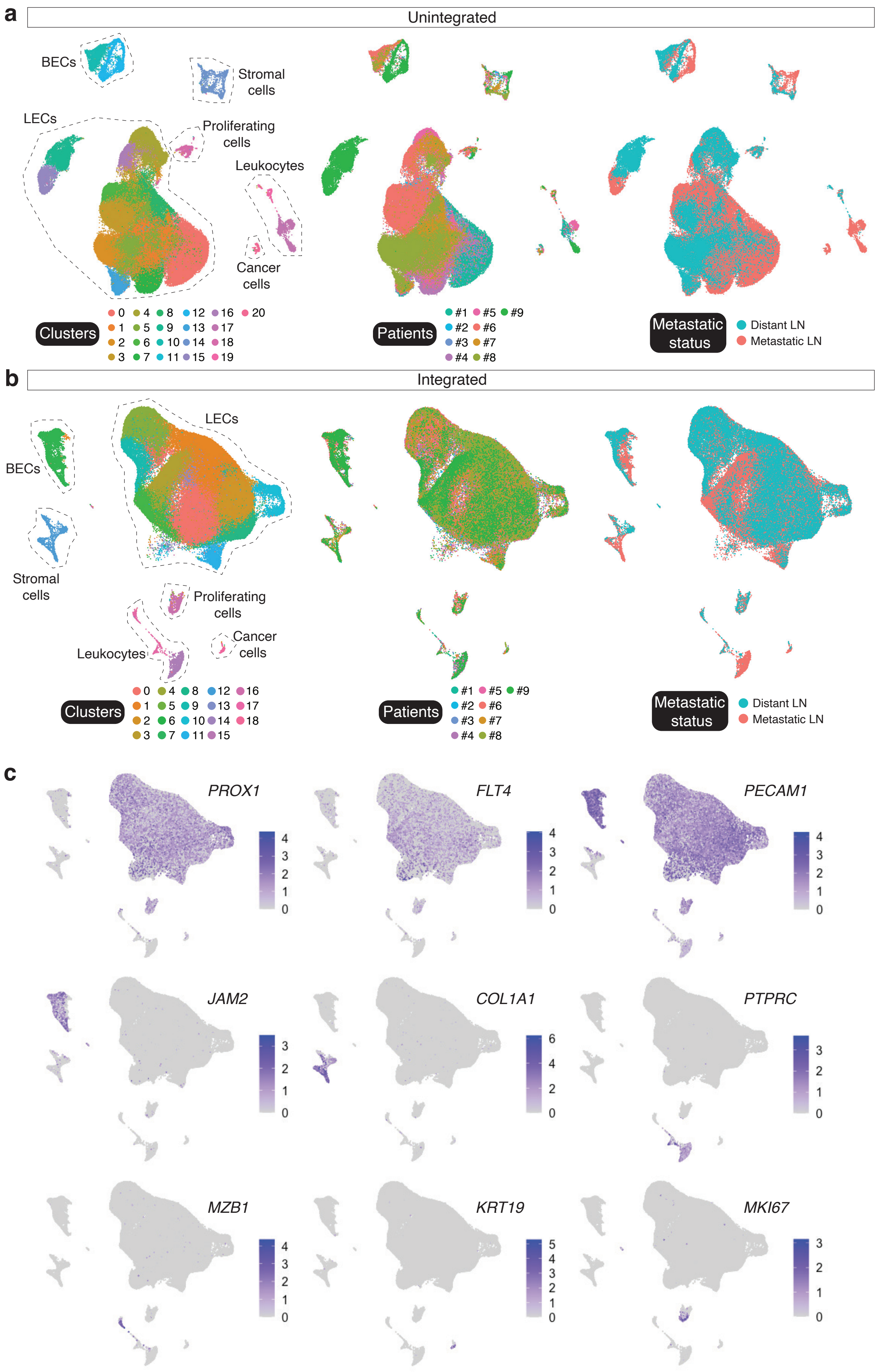


**Supplemental Figure 2. Cell subsets in LEC-enriched populations from metastatic and distant LNs. a, b,** UMAP plots of unintegrated (a) and integrated (b) LEC-enriched populations from distant and metastatic human LNs from 9 patient samples. Plots are colored by clusters (left), patients (middle), and metastatic state (right). **c,** Feature plots showing markers for each population. Markers include *PROX1* and *FLT4* for LECs, *JAM2* for BECs, *PECAM1* for ECs, *COL1A1* for stromal cells, *PTPRC* (CD45) for leukocytes, *MZB1* for plasmablasts, *KRT19* for cancer cells, and *MKI67* for proliferating cells.

#### Supplemental Figure 3

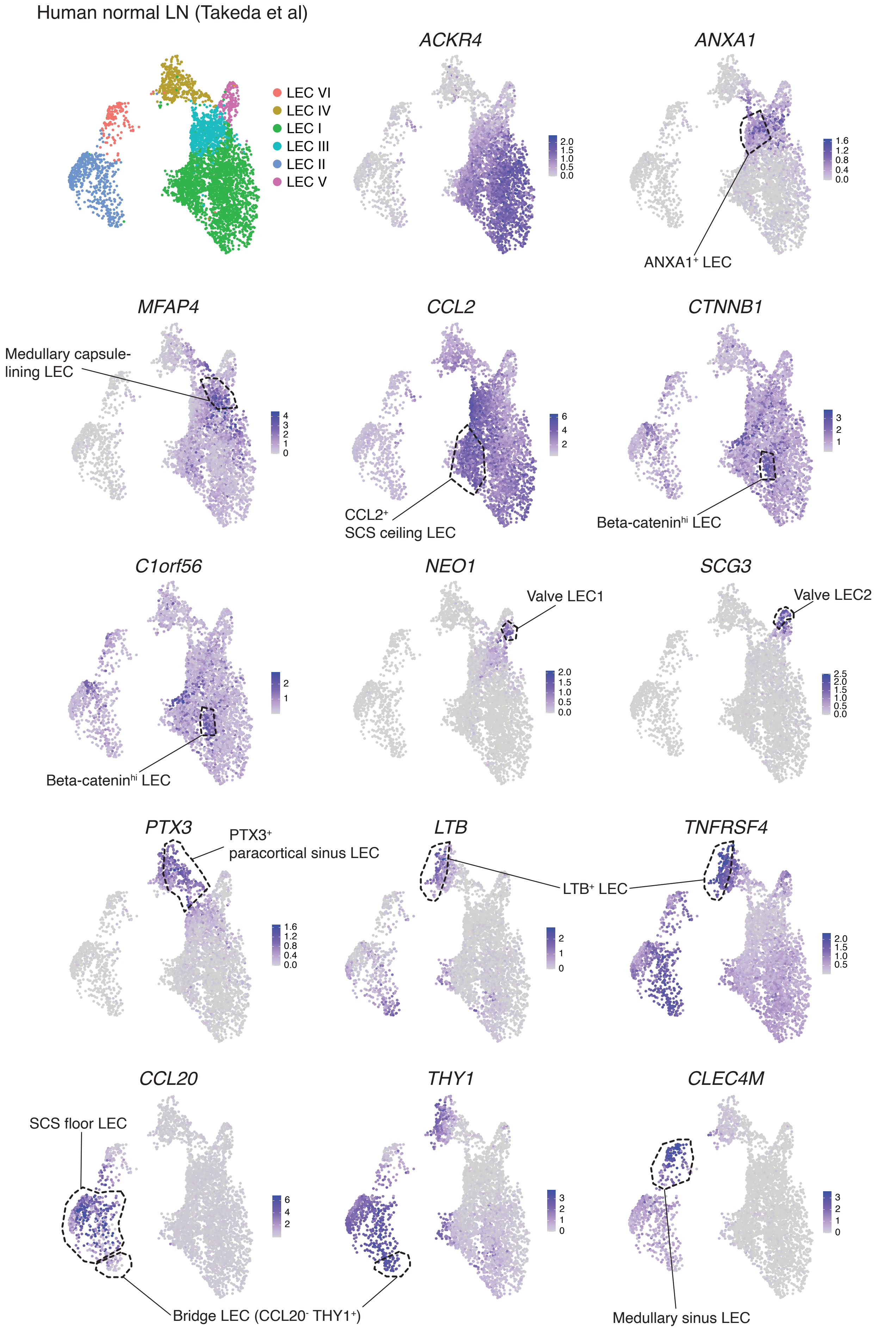


**Supplemental Figure 3. Validation of newly identified LEC subsets in a published dataset.** Reanalysis of our previously published human LN LEC scRNA-seq dataset (Takeda A et al., Immunity, 2019) to validate the newly identified LEC subsets. *ANXA1*$^{+}$ and *MFAP4*$^{+}$ medullary capsule-lining LECs were previously categorized as LEC III. *CCL2*$^{+}$ SCS ceiling LECs and *CTNNB1*$^{+}$ *C1orf56*$^{+}$ beta-catenin$^{hi}$ LECs are found within the LEC I fraction. *PTX3*$^{+}$ paracortical sinus LECs and *LTB*$^{+}$ *TNFRSF4*$^{+}$ LECs are within LEC IV. *CCL20*$^{+}$ SCS floor LECs and *CCL20*$^{-}$*THY1*$^{+}$ bridge LECs are within the LEC II fraction.

### Supplemental Figure 4

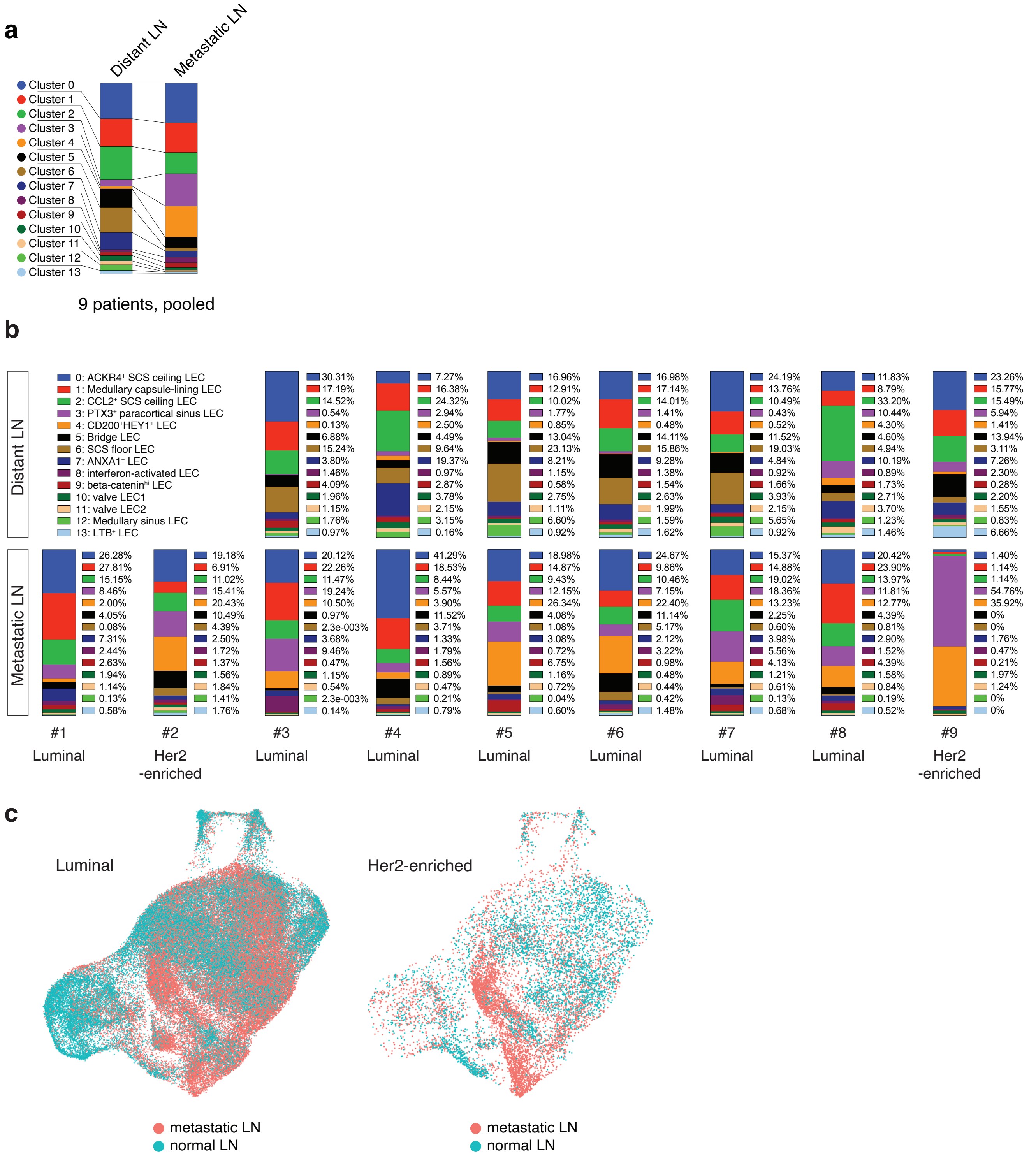


**Supplemental Figure 4. LN LEC subsets in luminal and Her2-enriched breast cancer patients. a, b,** Frequencies of each LEC subset in distant and metastatic LNs, presented as pooled data (a) and by individual patient (b). **c,** UMAP plots of LN LECs from luminal (left) and Her2-enriched (right) breast cancer patients.

#### Supplemental Figure 5

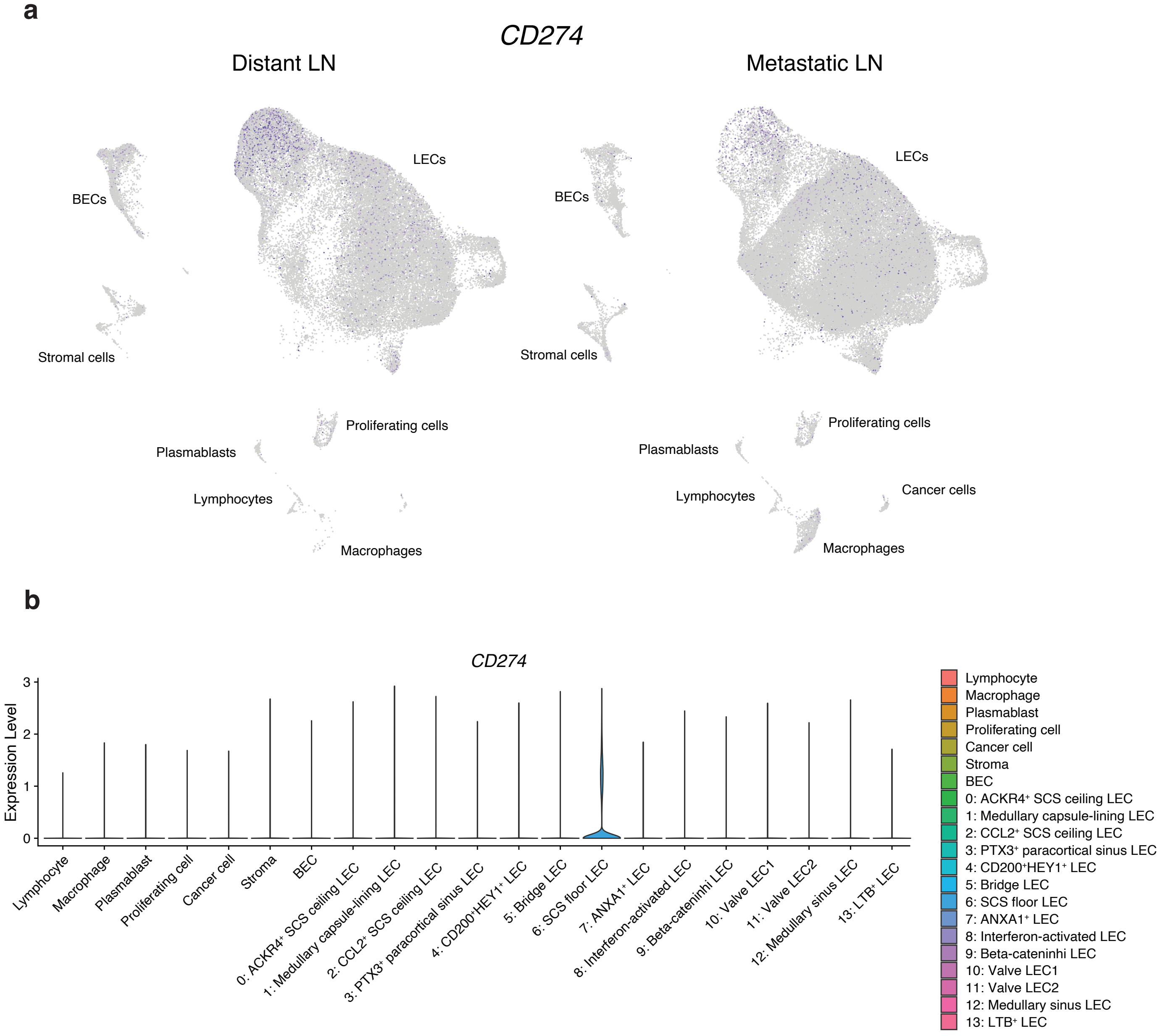


**Supplemental Figure 5. CD274 (PD-L1) expression in distant and metastatic LNs. a,** Feature plots showing *CD274* expression in distant and metastatic LNs. *CD274* was downregulated in metastatic LNs. **b,** Violin plot of *CD274* expression across all cell types in distant and metastatic LNs.

#### Supplemental Figure 6

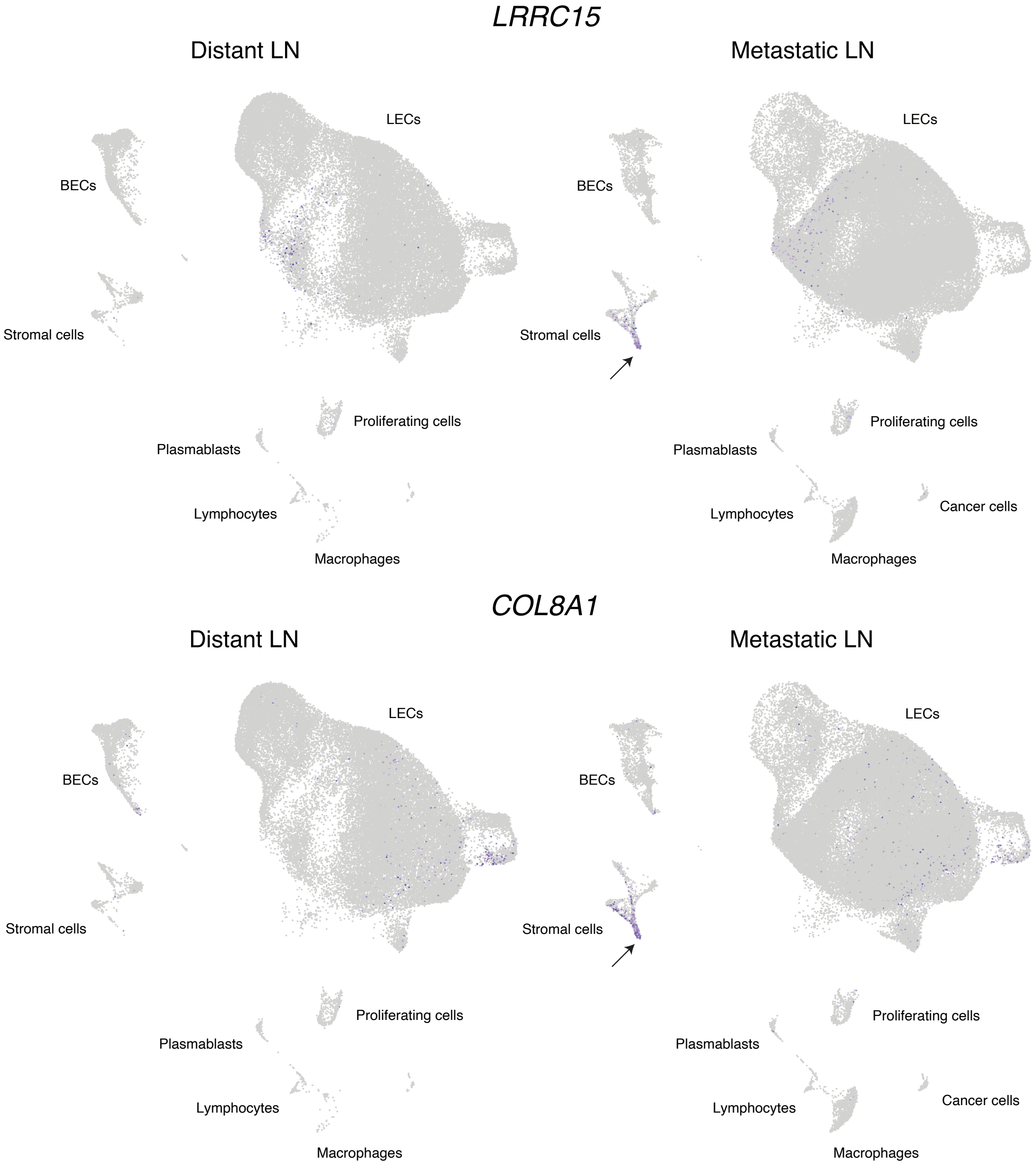


**Supplemental Figure 6. TGF-β-Dependent LRRC15⁺ Cancer-Associated Fibroblasts in Metastatic LNs.** LRRC15⁺ cancer-associated fibroblasts (indicated by arrows) are present in metastatic LNs but not in distant LNs. Feature plots display markers of LRRC15⁺ cancer-associated fibroblasts, including *LRRC15* and *COL8A1*.

### Supplemental Figure 7

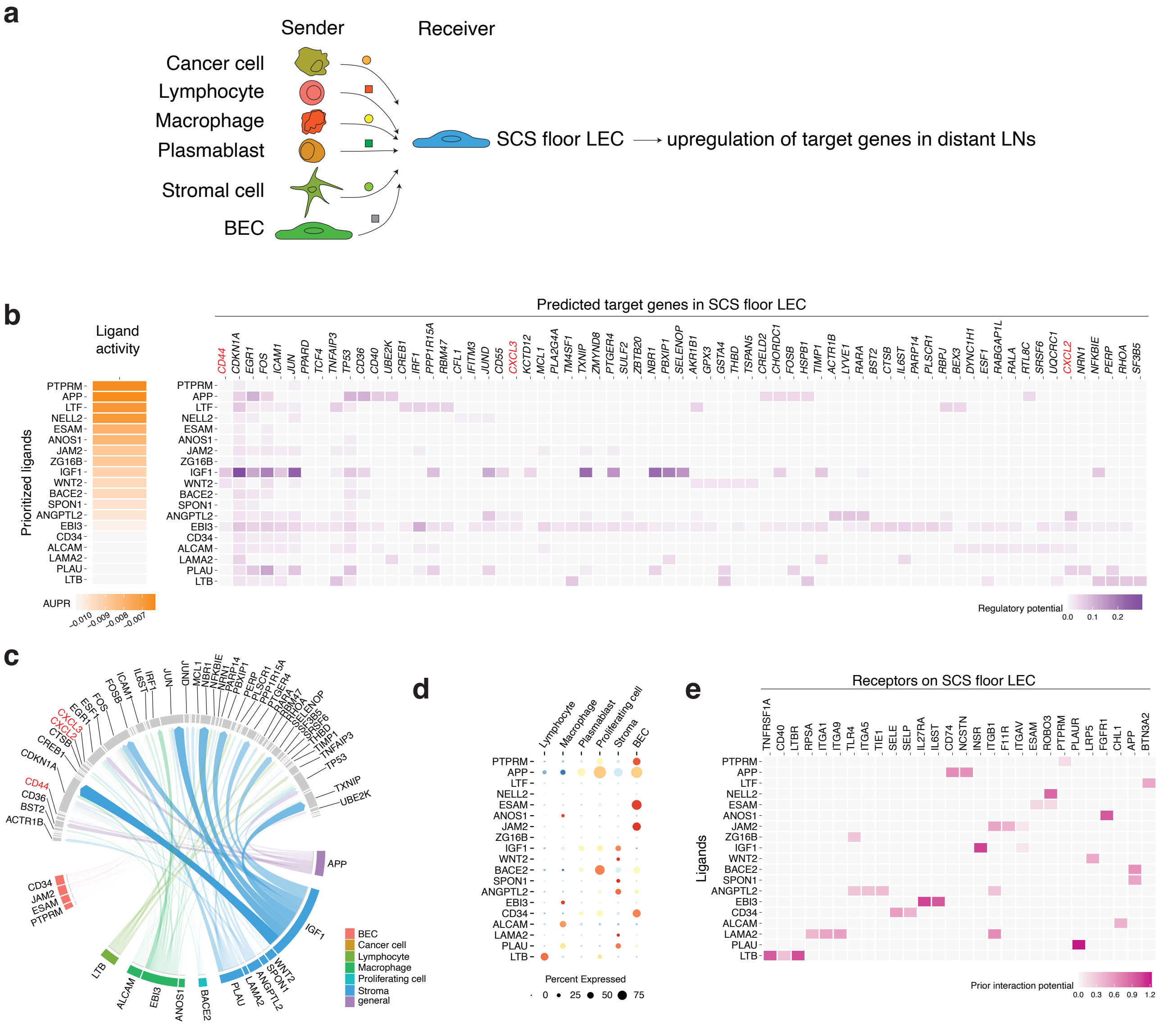


**Supplemental Figure 7. NicheNet Intercellular Communication Analysis of SCS Floor LECs. a,** Schematic illustration of the NicheNet analysis to investigate the mechanisms maintaining SCS floor LECs. SCS floor LECs are predominantly found in distant LNs, and genes upregulated in these LNs were considered as crucial for the maintenance of this LEC type. SCS floor LECs were designated as the receiver, while other LN cell types were set as senders. **b,** Predicted top ligands and their target genes in SCS floor LECs. **c,** Circos plot showing connections between predicted ligands from LN cells and their potential target genes in SCS floor LECs. **d,** Dot plot illustrating the expression of ligands in LN cells. **e,** Potential receptors expressed by SCS floor LECs associated with each predicted ligand.

**Supplemental Figure 8**

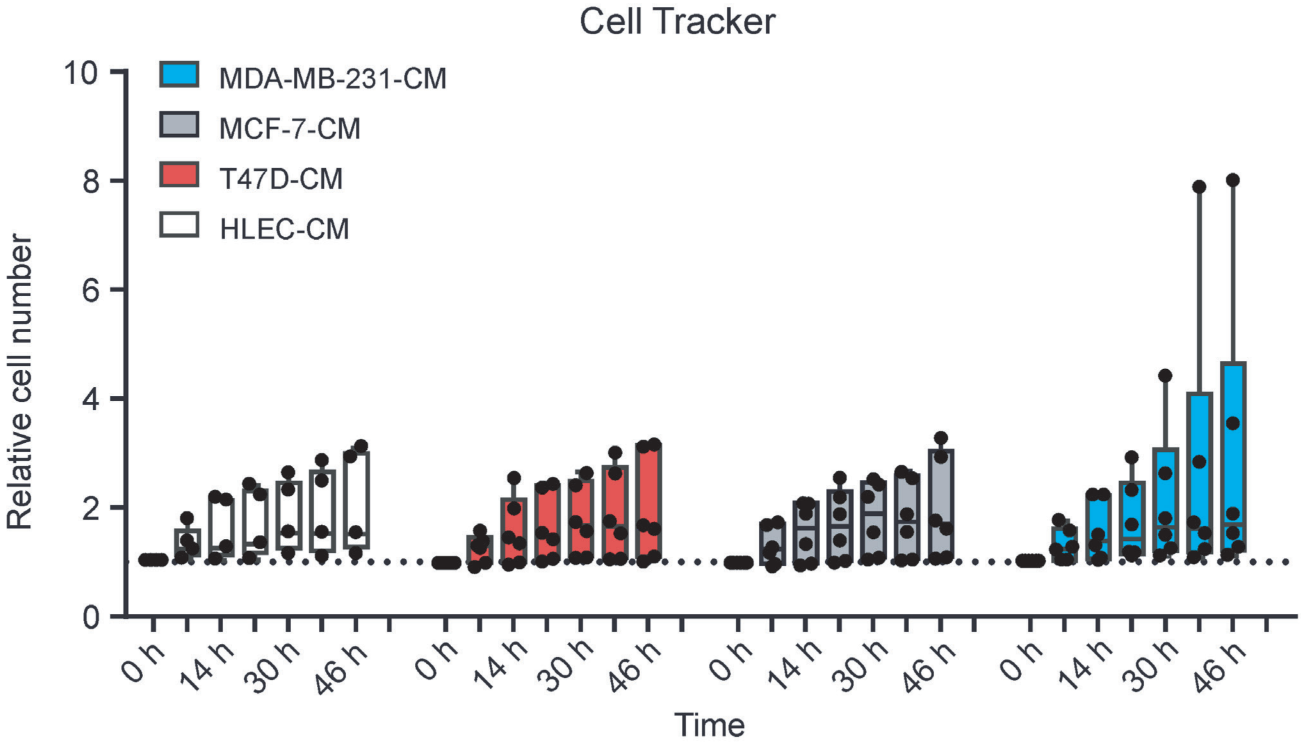


**Supplemental Figure 8. LEC proliferation in the presence of cancer cell CM.** The proliferation of LECs exposed to different CM is shown. LECs were labelled with 6.5 uM CellTracker Red, exposed to different CM and their relative cell number over the course of two days was determined at different timepoints. Data are shown as Tukey box plots (n=5-6). The center line of the box plots represents the median, the box the 25th to 75th percentiles and the whiskers inner fences.

#### Supplemental Figure 9

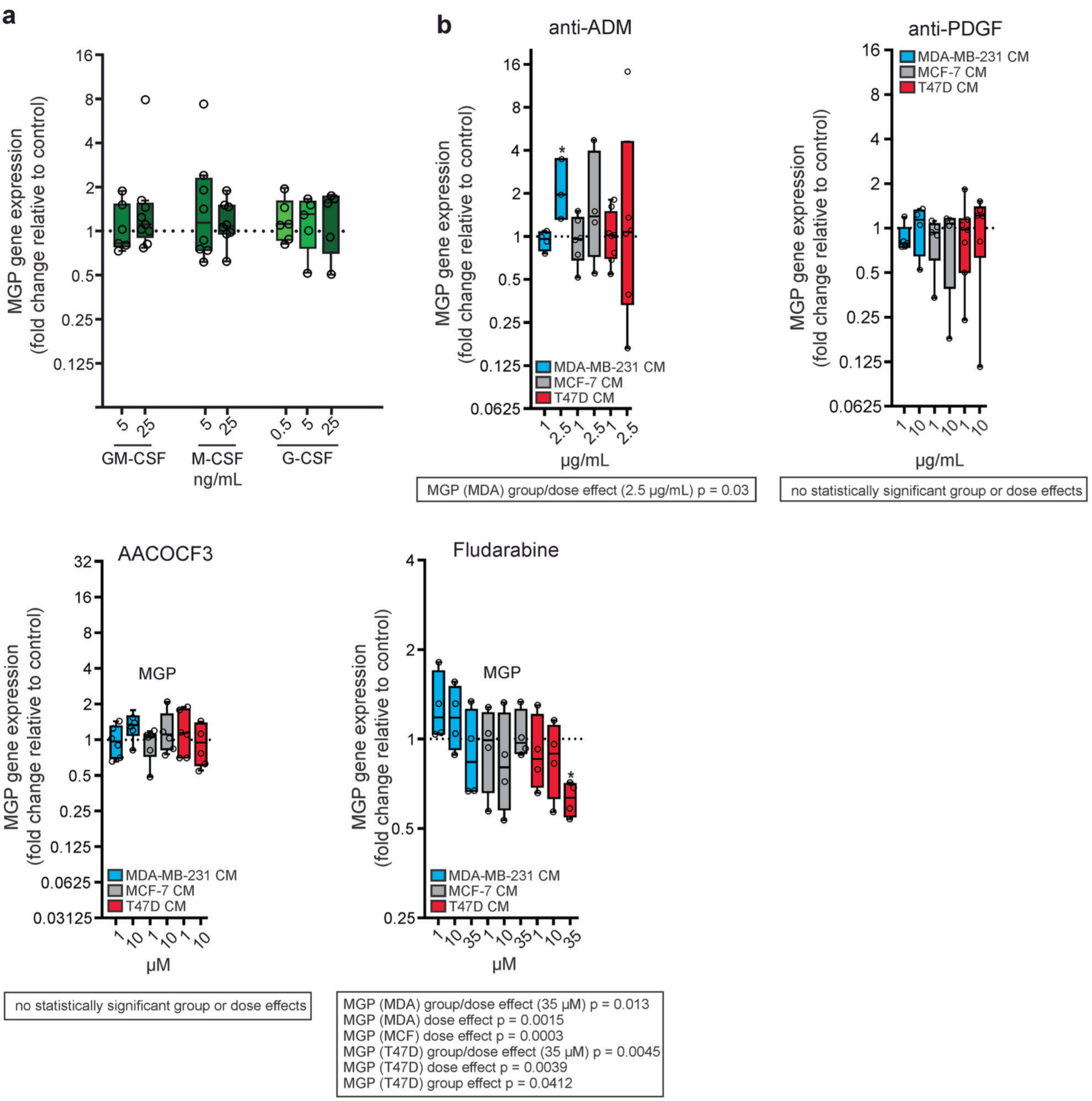


**Supplemental Figure 9. Gene expression of MGP after recombinant cytokine and antibody exposure. a,** Gene expression of MGP in LECs after direct exposure to recombinant GM-CSF (5 or 25 ng/mL, n=6-8), M-CSF (5 or 25 ng/mL, n=7-8) or G-CSF (0.5, 5 or 25 ng/mL, n=5-6), respectively are shown. Data are depicted as Tukey box plots. **b,** Gene expression of MGP in LECs are shown following their exposure to different modified CM. CM were generated in the presence of 1 and 2.5 µg/mL anti-ADM antibody, 1 and 10 µg/mL anti-PDGF antibody, 1 and 10 µM AACOCF3 and 1, 10 and 35 µM Fludarabine or their control antibody/substance, respectively. Data are shown as Tukey box plots (n=3-8). Data were analyzed with a linear mixed model. The center line of the box plots represents the median, the box the 25th to 75th percentiles and the whiskers inner fences. (*$p < 0.05$). Statistics of group and dose effects are presented within the boxes; differences in comparison to the controls (defined as 1) are indicated by stars.

**Supplemental Figure 10**

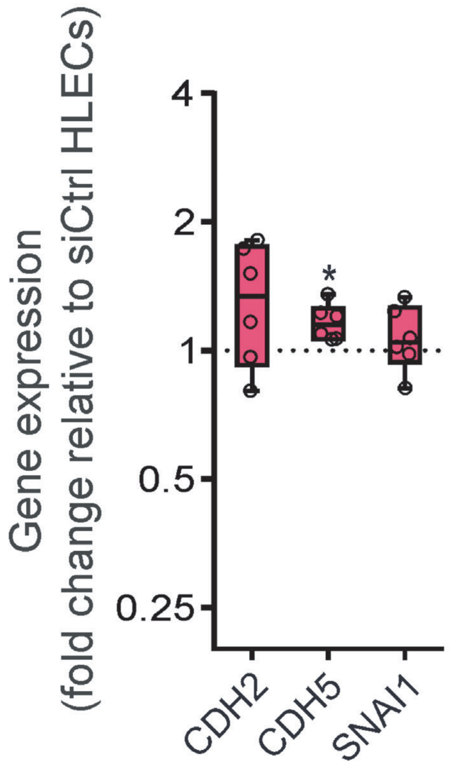


**Supplemental Figure 10. siMGP effect on EndMT.** Gene expression of CDH2, CDHS and SNAIl in MGP silenced LECs are shown as Tukey box plots (n=6) and analyzed with the Wilcoxon matched-pairs signed rank test. The center line of the box plots represents the median, the box the 25th to 75th percentiles and the whiskers inner fences.
